# A CRISPR/Cas9 genetically engineered organoid biobank reveals essential host factors for coronaviruses

**DOI:** 10.1101/2021.05.20.444952

**Authors:** Joep Beumer, Maarten H. Geurts, Mart M. Lamers, Jens Puschhof, Jingshu Zhang, Jelte van der Vaart, Anna Z. Mykytyn, Tim I. Breugem, Samra Riesebosch, Debby Schipper, Petra B. van den Doel, Wim de Lau, Cayetano Pleguezuelos-Manzano, Georg Busslinger, Bart L. Haagmans, Hans Clevers

**Affiliations:** Oncode Institute, Hubrecht Institute, Royal Netherlands Academy of Arts and Sciences and University Medical Center, Utrecht, Netherlands; Viroscience Department, Erasmus Medical Center, Rotterdam, Netherlands

## Abstract

Rapid identification of host genes essential for virus replication may expedite the generation of therapeutic interventions. Genetic screens are often performed in transformed cell lines that poorly represent viral target cells in vivo, leading to discoveries that may not be translated to the clinic. Intestinal organoids (IOs) are increasingly used to model human disease and are amenable to genetic engineering. To discern which host factors are reliable anti-coronavirus therapeutic targets, we generate mutant clonal IOs for 19 host genes previously implicated in coronavirus biology. We verify ACE2 and DPP4 as entry receptors for SARS-CoV/SARS-CoV-2 and MERS-CoV respectively. SARS-CoV-2 replication in IOs does not require the endosomal Cathepsin B/L proteases, but specifically depends on the cell surface protease TMPRSS2. Other TMPRSS family members were not essential. The newly emerging coronavirus variant B.1.1.7, as well as SARS-CoV and MERS-CoV similarly depended on TMPRSS2. These findings underscore the relevance of non-transformed human models for coronavirus research, identify TMPRSS2 as an attractive pan-coronavirus therapeutic target, and demonstrate that an organoid knockout biobank is a valuable tool to investigate the biology of current and future emerging coronaviruses.

## Main

Three highly pathogenic coronaviruses have spread to humans in the past two decades. The Severe Acute Respiratory syndrome coronavirus (SARS-CoV) emerged in 2002 and rapidly spread between continents, but was controlled by public health interventions. Middle East respiratory syndrome virus (MERS-CoV) -discovered in 2012-causes an ongoing outbreak in the Middle East with a high case-fatality ratio of 35%, but has not attained efficient human-to-human transmission. The latest, SARS-CoV-2, emerged at the end of 2019 and is the causative agent of the COVID-19 pandemic^1^. While vaccine development has taken off at a tremendous pace, drugs that target either the virus or host factors essential for virus replication have been more difficult to develop as this requires a deep understanding of coronavirus biology.

The first step in coronavirus replication is the attachment to host cells, which is dependent on the viral spike glycoprotein^2^. Although (proteo)glycans are often involved in the initial attachment, most coronavirus spikes require a specific transmembrane protein for entry. After receptor engagement, the next step in viral entry involves proteolytic cleavage of the spike protein. This cleavage step is performed by host proteases and destabilizes the spike, causing a conformational change and the subsequent fusion of viral and host membranes. This releases the viral ribonucleoprotein complex into a host cell and initiates replication.

Most of what we know on coronavirus cell biology stems from studies on 2D transformed cell lines such as the Vero E6 kidney cell line, derived from an African green monkey^3^. Work on cell lines has identified ACE2 as the entry receptor of SARS-CoV-2 and SARS-CoV, and DPP4 as the entry receptor of MERS-CoV^4–6^. Cell lines typically consist of a homogeneous population of poorly differentiated cells, potentially limiting the translatability of findings. As a case in point, chloroquine, an endocytosis inhibitor, has been proposed as a SARS-CoV-2 antiviral drug as it blocks SARS-CoV-2 entry in several cell lines^7^, yet clinical studies have failed to demonstrate efficacy in COVID-19 patients^8^. Along these lines, inhibiting protease groups in cell lines with relatively broad-acting inhibitors have revealed that spike protein cleavage can occur at the cell surface by transmembrane serine protease (e.g. TMPRSS family members) or in the endosome by cathepsins (e.g. Cathepsin B or L), depending on the cell line used. Recently published host gene loss-of-function screens in 2D cell lines have supported a role for Cathepsin L, but not TMPRSS2, in viral entry into VeroE6 cells, while the opposite was observed in a small scale CRISPR screen Calu-3 cells^9,10^. In addition, we and others have recently shown that in primary airway cells serine protease-(but not cathepsin-) inhibitors block viral entry^4,11^, but these inhibitors target all TMPRSS family members. Thus, it remains unknown whether in primary cells TMPRSS2 would be a realistic therapeutic target, or whether other TMPRSS family members could compensate for the loss of TMPRSS2. Similarly, several new host factors have recently been found to play a role in the SARS-CoV-2 replication cycle, such as NRP1 and NDST1, but it is unknown whether these genes could be used as anti-SARS-CoV-2 drug targets^12–14^.

There may exist significant differences between individual transformed cell lines, and between transformed and non-transformed cells in viral entry pathways. Intestinal organoid (IO) culture systems are an attractive platform to study virus-host interactions as they are amenable to CRISPR-Cas9 mediated gene editing to directly identify host proteins utilized by the virus. Their self-renewing nature offers an additional advantage: biobanks of characterized mutant IO clones can be established, stored and shared. Here we establish a biobank of mutant IOs in genes implied in coronavirus biology and test their role in coronavirus replication to discern which host factors may represent anti-coronavirus therapeutic targets. This biobank can be used as a tool to rapidly identify which genes are essential for virus entry when novel SARS-CoV-2 variants or novel zoonotic (corona)viruses emerge.

## Results

### Transcriptomic analysis of human IOs and airway cultures reveal conserved expression of coronavirus host factors

Multiple host factors such as entry receptors and proteases are involved in viral replication cycles^15^. Since organoids closely resemble the physiology of human tissues, we used IOs to assess the function of individual host factors that have been implicated in the SARS-CoV-2 replication cycle, or of other coronaviruses. We and others have previously shown that SARS-CoV-2 can replicate in human IOs^16–18^, consistent with observations of gastrointestinal symptoms in COVID-19^19,20^. Intestinal organoids are readily amenable to genetic engineering by CRISPR-Cas9^21^, allowing to test the role of host genes in the replication of SARS-CoV-2. We reasoned that individual host factors that upon loss-of-function affect coronavirus replication, represent interesting drug targets for the treatment of COVID-19. As spike protein cleavage is an essential step for viral entry, we focused on the proteases TMPRSS2, TMPRSS3, TMPRSS4, TMPRSS11D, TMPRSS13, Cathepsin B (CTSB), Cathepsin L (CTSL) and Furin, that have previously been implicated in the entry of SARS-CoV-2 or other coronaviruses^9,22,23^. Besides proteases, we included the following (putative) entry or attachment factors: the protease DPP4 (MERS-CoV), peptidase ANPEP (human coronavirus 229E receptor^24^, C-type lectin CLEC2B^25^, structural protein Vimentin^26^, glycoprotein CEACAM1^27^, tetraspanin CD9^28^, C-type lectin receptor CD209 (DC-SIGN)^29^, VEGF co-receptor NRP1^13,14^, MAVS which indirectly senses cytoplasmic RNA^30^, heatshock protein HSPA5^31^, the sulfotransferase NDST1^12^ and the RNA packaging protein ARC (upregulated upon SARS-CoV-2 infection^17^, Fig. S6E) as putative host factors involved in replication.

We first confirmed the expression of these host factors in human IOs using a previously generated dataset (Fig. 1A, S1A)^17^. To extend the expression analysis, we performed single cell RNA sequencing of expanding and differentiated IOs to assign host factors to individual cell types. The organoid atlas was supplemented by a recently published intestinal tissue dataset which we reanalyzed to allow for direct comparison^32^. Cells grouped into distinct stem cell, progenitor, goblet cell and enterocyte clusters (Fig. S1B-D). The proteases were ubiquitously expressed among the different lineages, with the exception of TMPRSS2 which displayed a slight bias towards enterocytes, and TMPRSS4 that was enriched in undifferentiated cells in both tissue and organoids (Fig. 1B). These findings contrasted with a recent study that used murine tissue data and identified goblet-cell enriched expression of TMPRSS2, while TMPRSS4 was mostly found in enterocytes^33^.

**Figure 1.**
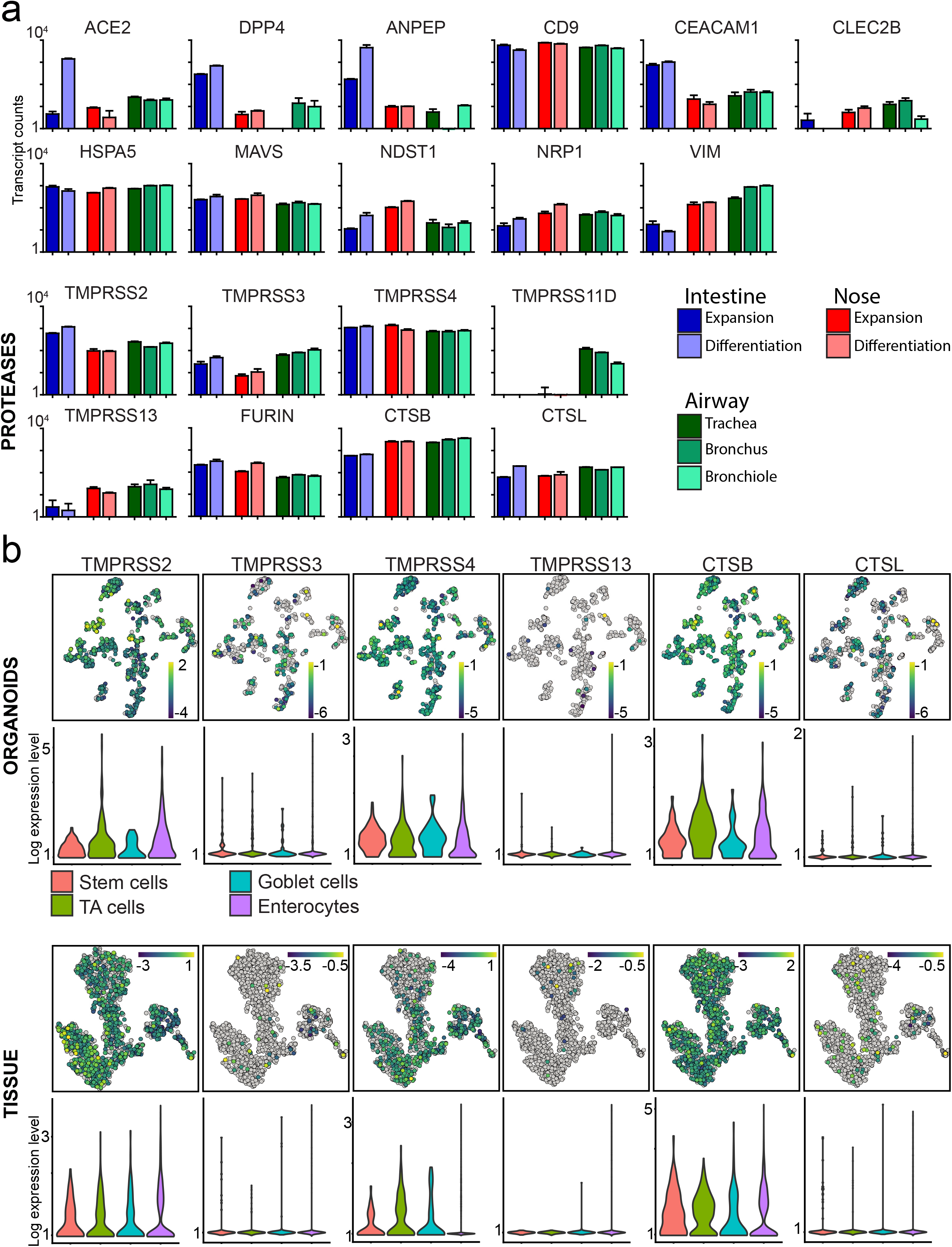
Expression levels of host genes potentially involved in SARS-CoV-2 biology in lung and intestinal organoids and tissue. a) Graphs depicting the transcript counts (logarithmic scale) determined by RNA sequencing of host factors and proteases in organoids derived from different parts of the airway and intestine. Airway organoids (trachea, bronchus and bronchiole) were differentiated as 2D ALI-cultures. b) t-SNE maps and violin plots displaying expression of host factors in the human intestinal organoid cell and human intestinal tissue atlases (in vivo atlas derived from^32^. Bars in t-SNE maps display color-coded normalized unique transcript expression (logarithmic scale).

We additionally performed RNA sequencing of 3D nasal organoids and 2D air-liquid interface differentiated airway organoids, confirming expression of all selected host genes in these epithelia, while relative expression levels varied (Fig. 1A, S1A, Table S1). Of note, the airway proteases TMPRSS11A and TMPRSS11D were expressed in lower airway cultures, while expressed to modest levels in IOs and nasal epithelial organoids (Fig. 1A, S1A).

### ACE2 is the obligate entry receptor of SARS-CoV and SARS-CoV-2

Having established expression of key coronavirus host factors, we generated an extensive biobank of IOs harboring a loss-of-function mutation in individual genes (Fig. 2A, S2–3). We employed transient transfection of a Cas9-EGFP encoding plasmid that included a site-specific guide RNA (sgRNA). Transfected cells were GFP-sorted to establish clonal lines (12-24 lines per gene), expanded and sequenced to identify loss-of-function clones. We successfully established 2-6 mutated clonal IO cultures for all genes with the exception of HSPA5 (0 lines) and Furin and ARC (1 line) (Fig. 2B, Table S2). HSPA5 is a heatshock protein that was recently identified as an essential gene for cell survival^34^.

**Figure 2.**
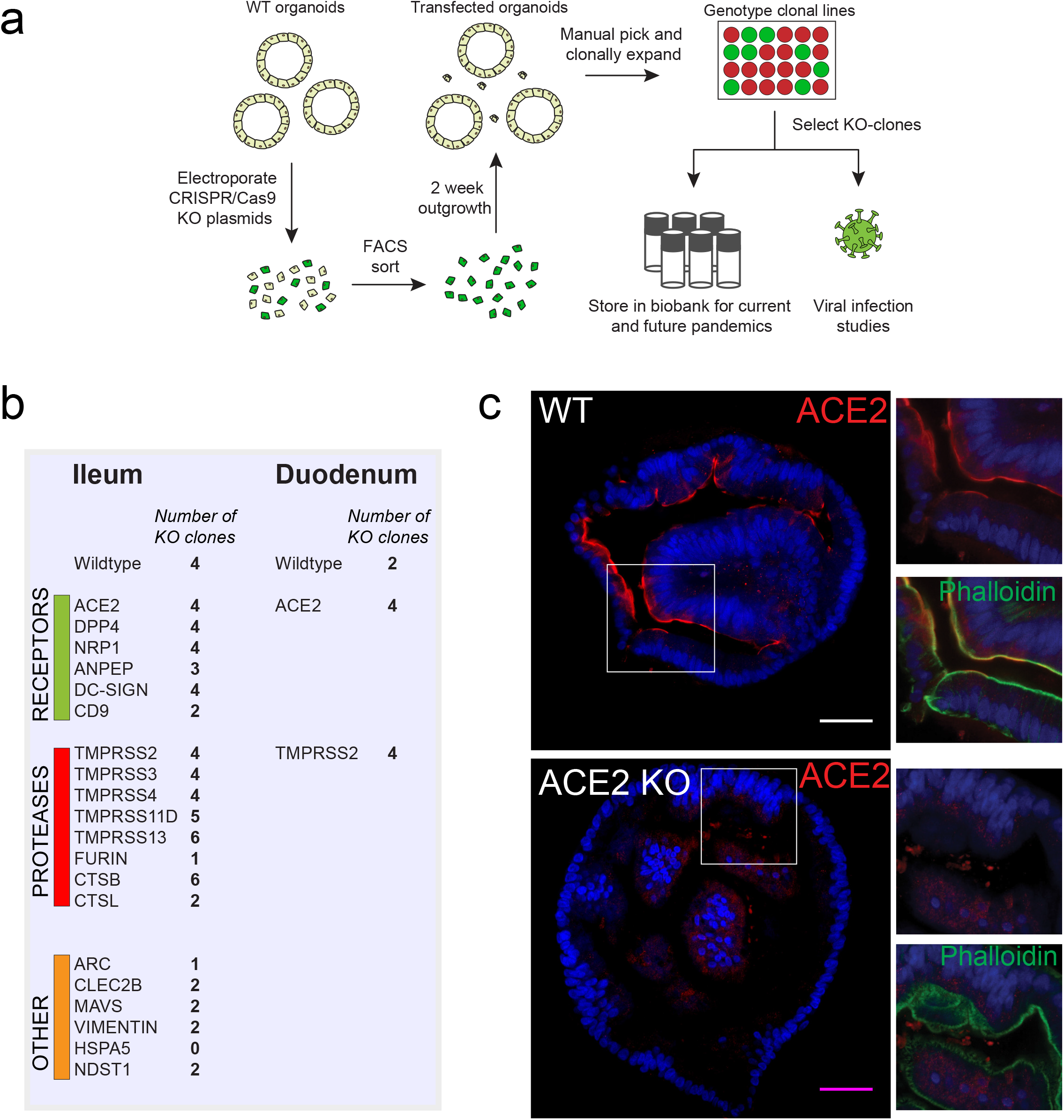
Generation of a coronavirus host gene knockout biobank. a) Overview of the workflow of generation of gene knockouts. b) Overview of the number of clones that were amplified and biobanked in this study. c) Immunofluorescent staining of WT and ACE2 KO organoids. ACE2 locates to the apical membrane and is absent in mutant organoids. Scale bars are 50 μm. Contrast was enhanced in unzoomed images for visualization purposes.

We first focused on ACE2, generally accepted to be the entry receptor for SARS-CoV-2 and SARS-CoV, as based on crystallographic evidence and overexpression studies^5^. Moreover, transgenic mice expressing human ACE2 are susceptible to infection with the virus^35^. Multiple additional (co-)receptors have been proposed for SARS-CoV-2, including CD209/DC-SIGN, and NRP1^13,14,36^. Recent findings in human IOs have found no correlation between infectability of cells and their levels of ACE2 expression, suggesting the potential existence of alternative entry receptors^37^. In line with these findings, we observed both ACE2-positive and ACE2-negative SARS-CoV-2 infected cells in IOs (Fig. S4), although absence of surface-ACE2 could also indicate downregulation after infection^37^ or reflect expression levels under the detection limit of immunofluorescence staining.

To unequivocally demonstrate that physiological levels of ACE2 are essential for SARS-CoV-2 entry into non-transformed human epithelial cells, we analyzed mutant ACE2 IOs for their ability to support SARS-CoV-2 replication. ACE2, located on the apical membrane of cells in wildtype IOs, was lost in mutant clones (Fig. 2C). ACE2-deficient IOs were fully resistant to SARS-CoV-2 infection (Fig. 3A). Indeed, we did not detect SARS-CoV-2 infected cells in ACE2-knockout organoids by immunofluorescence (Fig. 3B). Similarly, infection with SARS-CoV was abrogated in ACE2-knockout organoids (Fig. S5A). We concluded that ACE2 is the obligate entry receptor for SARS-CoV-2 and SARS-CoV, and that no redundancy exists with other surface proteins in intestinal epithelial cells. The presence of infected cells that appear ACE2-negative, implies either that surface receptors are downregulated upon infection or that low levels of ACE2 proteins suffice. To confirm that viral entry occurs through the apical membrane - where ACE2 is located - we attempted viral infection following our standard approach in which organoids are mechanically disrupted, and using intact organoids where only the basolateral surface is exposed. We could observe viral replication only in disrupted organoids, supporting an obligate apical entry route for SARS-CoV-2 (Fig. S5B).

**Figure 3.**
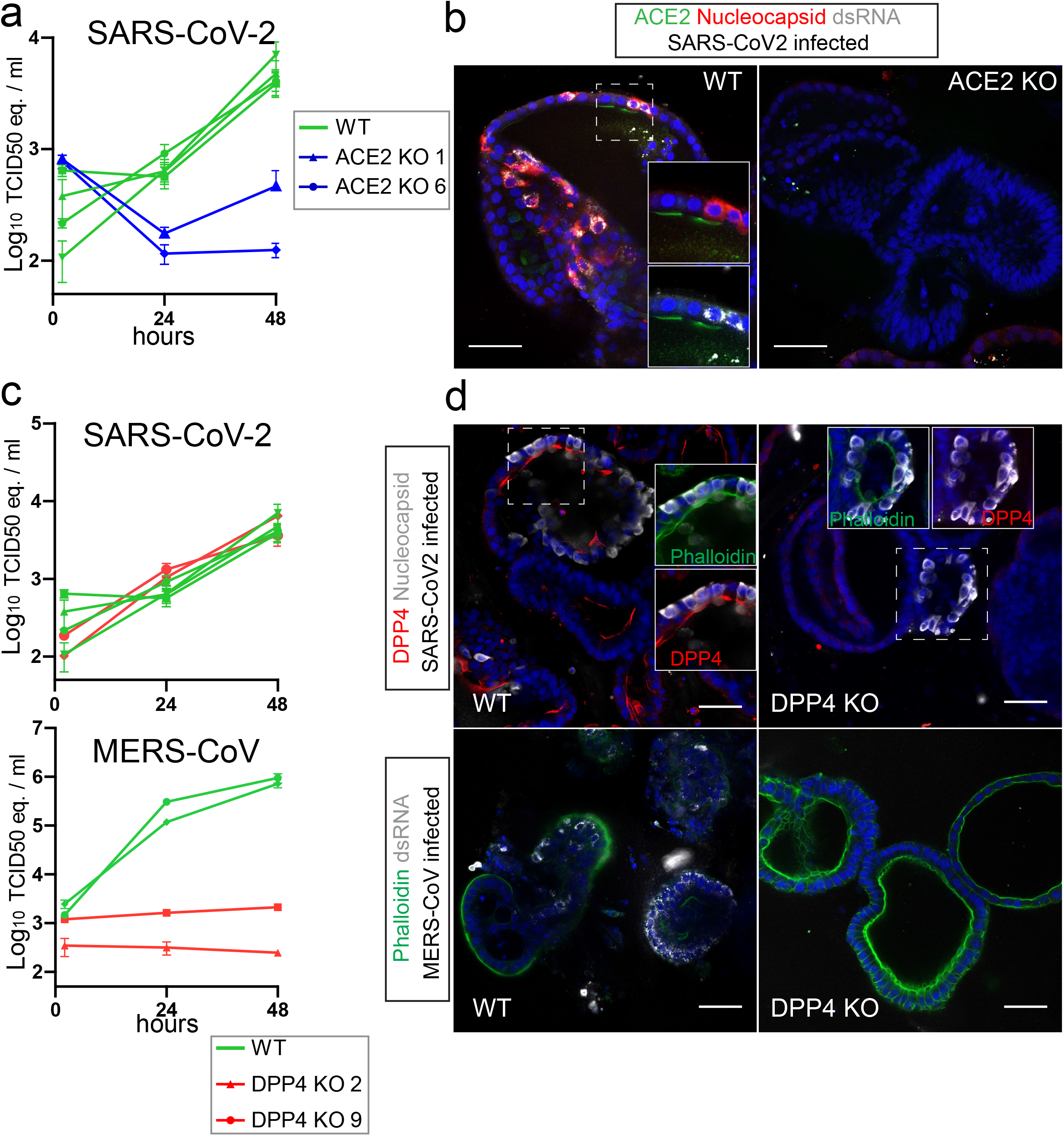
ACE2 and DPP4 are the obligate entry receptors for SARS-CoV-2 and MERS-CoV respectively. a) qPCR analysis targeting the E gene to quantify viral replication of wildtype (WT) and ACE2 knockout (KO) organoids. Error bars represent SEM. Each data point represents the mean of 3 replicates. b) Immunofluorescent staining of WT and ACE2 KO organoids, infected with SARS-CoV-2 visualized by nucleocapsid protein and dsRNA. ACE2 locates to the apical membrane and is absent in mutant organoids. c) qPCR analysis targeting the E gene (SARS-CoV-2) or upE region (MERS-CoV) to quantify viral replication in WT and DPP4 KO organoids. Each data point represents the mean of 3 replicates. d) Immunofluorescent staining of MERS-CoV (visualized by dsRNA) and SARS-CoV-2 (visualized by nucleocapsid protein) infected organoids. DPP4 locates to the apical membrane and is absent in mutant organoids. Scale bars are 50 μm.

### MERS-CoV infects human IOs in a DPP4-dependent manner

DPP4 has been shown to be the entry receptor for MERS-CoV by spike co-immunoprecipitation and overexpression in non-susceptible cells^6^. We first established that human IOs allow replication of MERS-CoV (Fig. S6A). In contrast to SARS-CoV-2, MERS-CoV caused extensive cell death, killing the majority of cells in organoids within 48 hours of infection (Fig. S6A). Transcriptomic analysis revealed a strong upregulation of heat shock- and unfolded protein responses, while interferons were effectively repressed (Fig. S6B-E, Tables S3-4). This was consistent with previous reports that MERS-CoV encodes an extensive set of proteins that inhibit interferon responses^38^. We generated loss-of-function DPP4-mutant IO clones (Table S2). An infection assay on two of these clones revealed that MERS-CoV replication was fully blocked, while SARS-CoV-2 replicated in the DPP4-mutant organoids at control levels (Fig. 3C). Immunofluorescence confirmed loss of DPP4 protein in mutant clones as well as the successful replication of SARS-CoV-2 -but not of MERS-CoV-in these organoids (Fig. 3D). Conversely, MERS-CoV propagated in ACE2-deficient organoids at control levels (Fig. S5A).

### Loss-of-function screen of host proteases reveals essential role in viral replication for TMPRSS2 but not other TMPRSS family members or Cathepsins

We next analyzed all IO lines that were mutant in proteases for their ability to support SARS-CoV-2 replication. Knockout of TMPRSS2 effectively blocked viral replication, while mutation of any of other TMPRSS-genes had no effect (Fig. 4A-B). Complete depletion of TMPRSS2 protein was confirmed using immunohistochemistry (Fig. S7A). These experiments were performed using a VeroE6-propagated stock and recent work has pointed out that propagation on VeroE6 cells can lead to culture adaptive mutations in the multibasic cleavage site^39–43^. The VeroE6 stock used in this study was deep-sequenced^39^ and was 64.2% wild-type in the RRAR (spike positions 682-685) multibasic cleavage site. We detected another mutation adjacent to the multibasic cleavage site (S686G) at a frequency of 45.4%. Viruses with multibasic cleavage site cleavage site mutations, including S686G, were shown to slightly increase cathepsin usage by ~20%^39^, indicating that the majority of these viruses still used serine protease-mediated entry.

**Figure 4.**
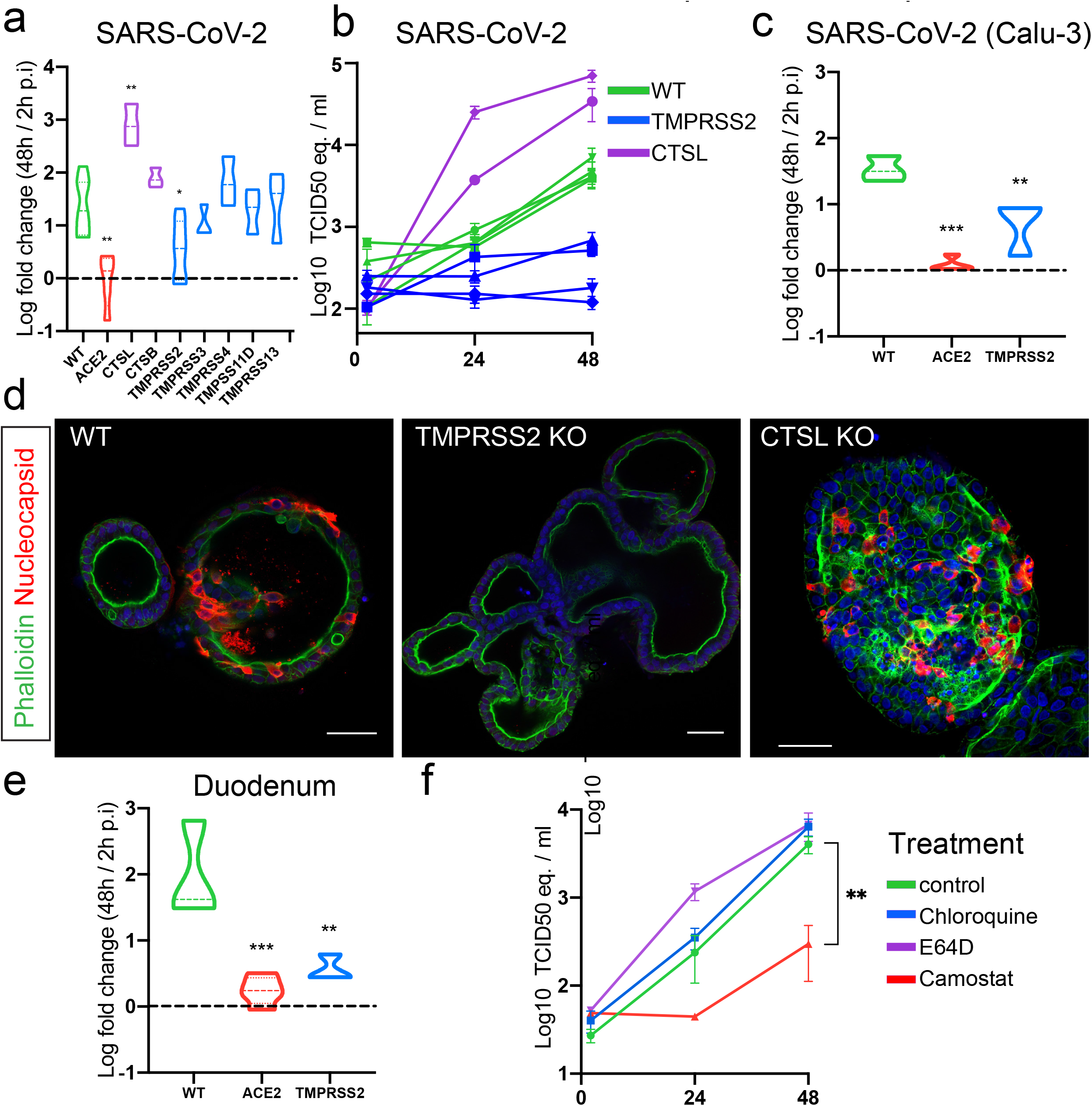
Loss-of-function screen of host proteases reveals essential role for TMPRSS2 but not Cathepsins in viral replication. a) Violin plot displaying the Log10 ratio between the viral titer at 48 hours compared to 2 hours post infection (p.i.) quantified by qPCR targeting the E gene. The dotted line indicates a fold change of 1. Dotted lines within the violins indicate the median and quartiles. Data from replication curves in Fig. 3A and 3C is included. N ≥ 3. b) qPCR analysis targeting the E gene to quantify viral replication of wildtype organoids and cells harboring a loss-of-function of a single host factor. Error bars represent SEM. Each data point represents the mean of 3 replicates. c) Violin plot displaying the Log10 ratio between the viral titer at 48 hours compared to 2 hours post infection (p.i.) quantified by qPCR targeting the E gene in organoids infected with SARS-CoV-2 propagated on Calu-3 cells. The dotted line indicates a fold change of 1. Dotted lines within the violins indicate the median and quartiles. N ≥ 3. d) Immunofluorescent staining of SARS-CoV-2 infected WT and CTSL and TMPRSS2 KO organoids. Virus is visualized by nucleocapsid protein. TMPRSS2 KO organoids do not facilitate viral infection, although very rare, infected cells can be observed. CTSL KO organoids display abundant viral infection. Scale bars are 50 μm. e) Violin plot displaying the Log10 ratio between the viral titer at 48 hours compared to 2 hours post infection (p.i.) quantified by qPCR targeting the E gene in organoids of the human duodenum. The dotted line indicates a fold change of 1. Dotted lines within the violins indicate the median and quartiles. N ≥ 3. f) qPCR analysis targeting the E gene to quantify viral replication of organoids treated with the serine protease inhibitor Camostat, chloroquine or cysteine protease inhibitor E64D. Error bars represent SEM. Each data point represents the mean of 3 replicates. P < 0.05 *; P<0.01**; P<0.001***; P<0.0001****.

We have previously shown that propagation in TMPRSS2-expessing Calu-3 cells prevents culture adaptation. Using this Calu-3 stock that was completely non-adapted^39^, we confirmed the dependency of SARS-CoV-2 on TMPRSS2 (and ACE2) (Fig. 4C). Immunofluorescence of TMPRSS2-deficient organoids showed absence of viral spread (Fig. 4D). This implied that TMPRSS2 is the main proteolytic activator of the SARS-CoV-2 spike protein. In contrast to knockout screens in VeroE6 cells that showed that the endocytic pathway protease Cathepsin L was essential for SARS-CoV-2 entry^9^, SARS-CoV-2 replicated -if anything-more efficiently in Cathepsin L-mutant than in wildtype organoids (Fig. 4A-B). Efficient depletion of Cathepsin L was supported using western blot analysis (Fig. S7B). We confirmed the obligate role of TMPRSS2 for SARS-CoV-2 replication in IOs derived from a different donor and from another segment (duodenum) of the human intestine (Fig. 4E).

Since we observed differential expression of multiple proteases - including upregulation of cathepsins - in differentiated organoids, we additionally assessed TMPRSS2-dependency in differentiated intestinal cells (Fig. S7C). After 5 days of differentiation, both wildtype and TMPRSS2-mutant organoids were infected with SARS-CoV-2. SARS-CoV-2 replicated efficiently in differentiated organoids, as we reported previously^17^. Viral replication was greatly diminished in TMPRSS2-deficient organoids, suggesting dependency on this protease across different intestinal cell types (Fig. S7D)

To assess redundancy in single TMPRSS- or cathepsin-mutant organoids, we additionally generated organoids mutant for both CTSL/CTSB, or TMPRSS2/4, the most abundantly expressed cathepsins and serine proteases in the intestinal epithelium. Previous work implied a role for TMPRSS4 in viral entry in the intestinal epithelium^33^. We did not observe reduced replication when both cathepsins were lost. Moreover, TMPRSS4 knock-out in a TMPRSS2-mutant background did not further decrease infectivity (Fig. S7E). In line with this, the broad serine protease inhibitor camostat did not affect replication in TMPRSS2-deficient IOs (Fig. S7F). We concluded that the cathepsins and TMPRSS4 do not play a role in viral entry in the intestinal epithelium.

To confirm that the endocytic pathway is dispensable for viral entry, we treated IOs with 1) the endosomal pathway inhibitor chloroquine, the cathepsin protease inhibitor E64D, or the broad serine protease inhibitor camostat. These drugs were well-tolerated with no growth impairment at the concentrations used (Fig. S8A-B). As published previously, chloroquine was effective in VeroE6 cells (Fig. S8C)^7^. While camostat effectively inhibited viral replication, chloroquine and E64D did not affect replication in IOs (Fig. 4F). E64D-treated organoids displayed a trend towards more efficient viral replication (Fig. 4F), consistent with observations in the Cathepsin L-mutant organoids (Fig. 4A-B). We concluded that Cathepsin-mediated entry through the endosomal route may be the central port of viral entry in cell lines, but not in IOs, in which SARS-CoV-2 enters through the activity of TMPRSS2 (Fig 4A-B). These observations may also explain why (hydro)-chloroquine has emerged from cell line studies but has proven ineffective in the clinic.

### Redundancy for non-protease host factors in viral replication

We further tested IOs mutant in a range of non-protease host factors to assess their role in coronavirus replication, of which some have already been linked to the SARS-CoV-2 replication cycle. NRP1 recently attracted attention as a novel co-receptor for SARS-CoV-2 in two separate studies that used Hela, HEK293T and the colorectal cancer cell line Caco-2^13,14^. These findings were substantiated by x-ray crystallography data supporting binding of the viral spike protein to NRP1 SARS-CoV-2 infection was significantly inhibited by NRP1-blocking antibodies^13,14^. Additionally, CD209 was recently identified as potential SARS-CoV-2 receptor, and facilitated viral entry in HEK-293 cells when overexpressed^36^. A recent study found that SARS-CoV-2 can bind heparan sulfate on the cell surface through its spike protein. When enzymes involved in the sulfation of heparan sulfate, including NDST1, were knocked out in Hep3B cells, viral replication was almost entirely abolished^12^. None of these, nor the additional host factors we assessed, significantly impacted on viral replication when mutated in IOs (Fig. 5A, Fig. S9). We conclude that all of these proteins would therefore not be viable drug targets for the treatment of COVID-19 (Fig. 5B). Further studies may assess whether loss of these factors influence the cellular response to coronaviruses in any other way than replication efficiency.

**Figure 5.**
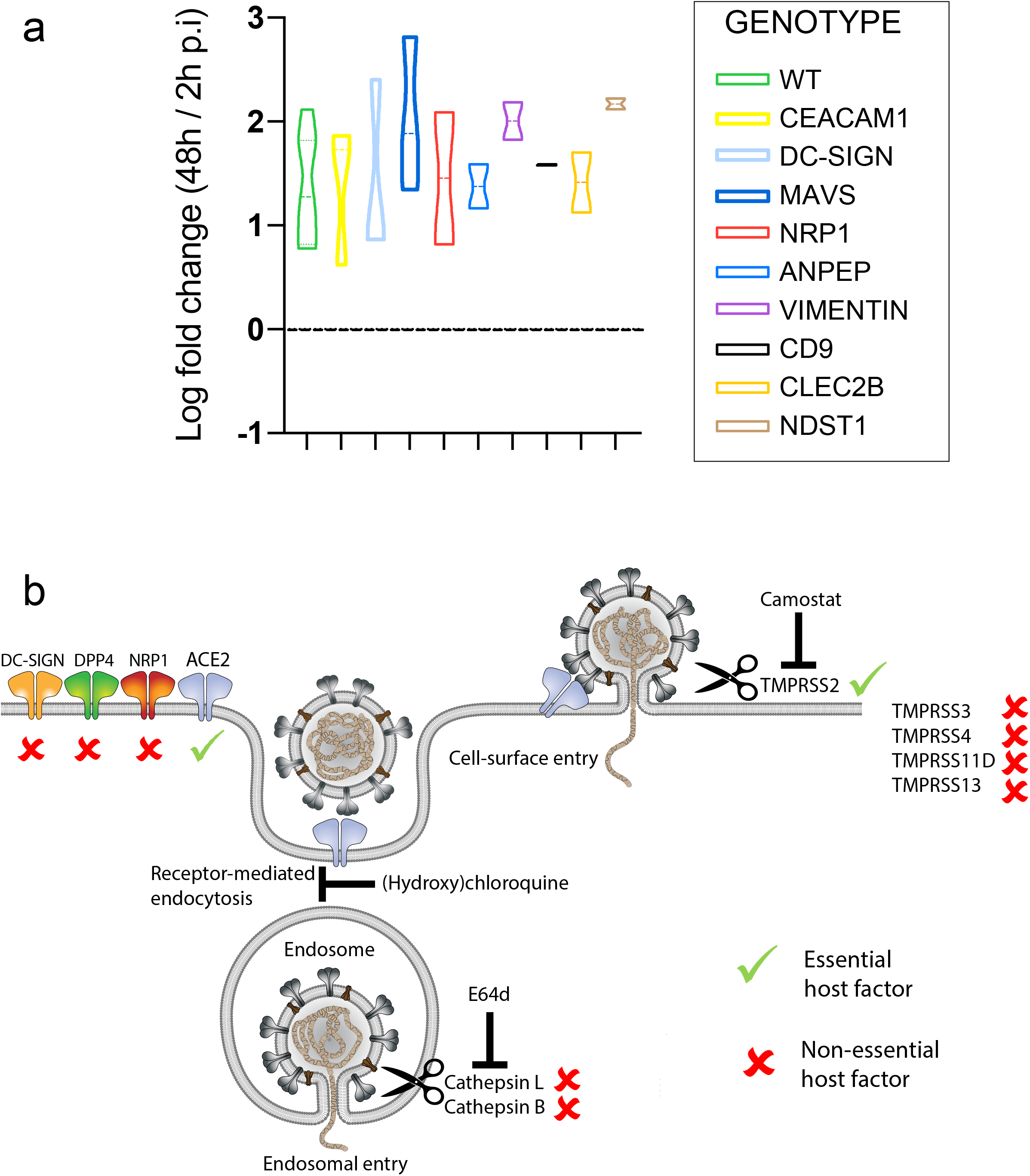
Replication of SARS-CoV-2 in additional proposed host factors. a) Violin plot displaying the Log10 ratio between the viral titer at 48 hours compared to 2 hours post infection (p.i.) quantified by qPCR targeting the E gene in wildtype organoids and cells harboring a loss-of-function of a single host factor. Graphs display the ratio between viral titer at 48 hours compared to 2 hours post infection (p.i.). The dotted line indicates a fold change of 1. Dotted lines within the violins indicate the median and quartiles. N ≥ 2. The WT data is redispayed from Fig. 4A. b) Model of host entry receptors and proteases involved in the entry of SARS-CoV-2. Essential and non-essential host factors based on the phenotypes in IO mutants are marked

### TMPRSS2-dependency in SARS-CoV-2 B.1.1.7, SARS-CoV and MERS-CoV

The mutant host factor KO biobank can readily be employed when new coronaviruses or viral strains appear, to assess the dependency on host factors and identify druggable targets. We first tested whether the same TMPRSS2-dependency exists for the other two coronaviruses. SARS-CoV replication was strongly diminished upon TMPRSS2 loss, while MERS-CoV replication was reduced more modestly (Fig. 6A). The latter potential redundancy may be explained by the presence of two functional multibasic cleavage sites in the MERS-CoV spike, whereas SARS-CoV-2 and SARS-CoV possess one and none, respectively^22^. Both viruses could replicate in the absence of Cathepsin L, suggesting that coronaviruses generally do not use the endosomal entry route in primary epithelial cells as present in organoids (Fig. 6A).

**Figure 6.**
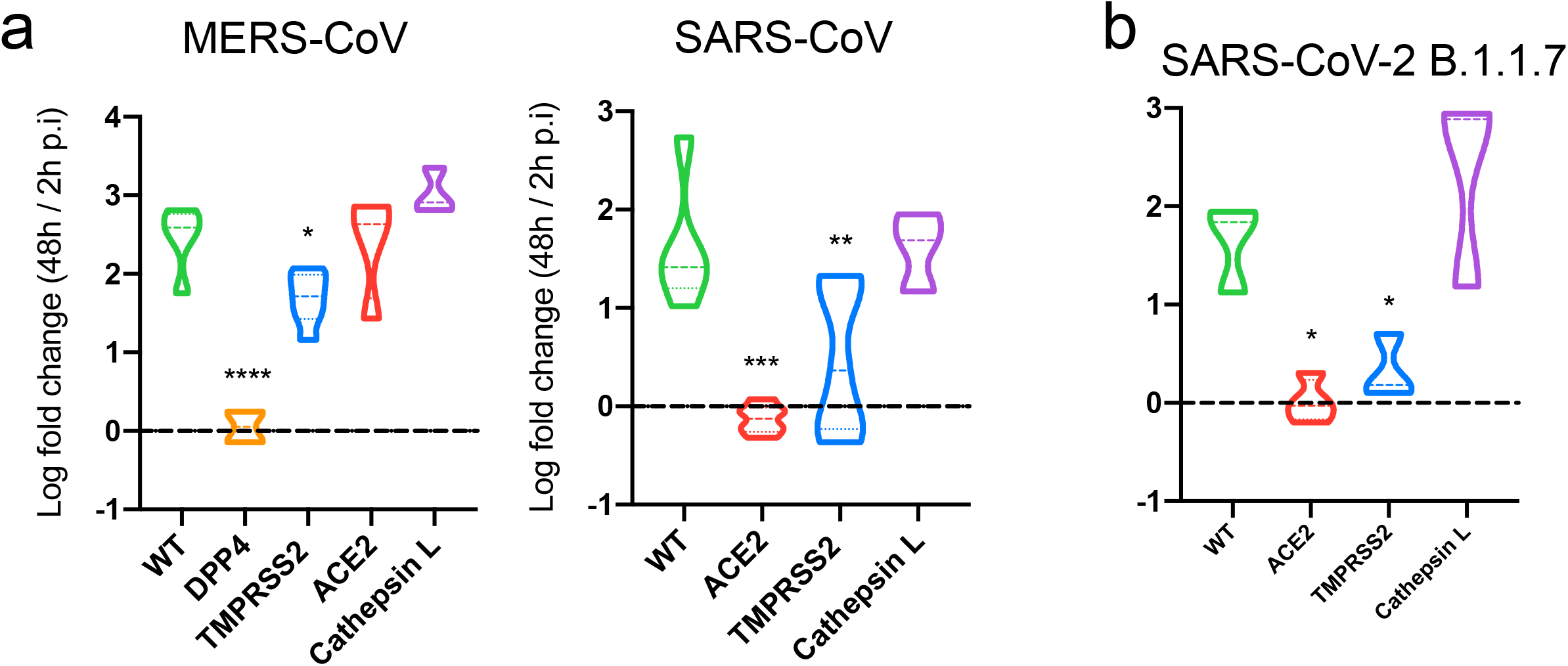
Host protease dependency in the SARS-CoV-2 strain B.1.1.7 and other coronaviruses. a) Violin plot displaying the Log10 ratio between the viral titer at 48 hours compared to 2 hours post infection (p.i.) quantified by qPCR targeting the the N gene (SARS-CoV) or upE region (MERS-CoV) to measure viral replication in WT and TMPRSS2 KO organoids. DPP4 and ACE2 data include replication curves in Fig. 3C and S4A. The dotted line indicates a fold change of 1. Dotted lines within the violins indicate the median and quartiles. N ≥ 3. b) Violin plot displaying the Log10 ratio between the viral titer at 48 hours compared to 2 hours post infection (p.i.) quantified by qPCR targeting the E gene to measure viral replication of the SARS-CoV-2 B.1.1.7 strain in intestinal organoid cells harboring a loss-of-function mutation in the TMPRSS2 and CTSL proteases, or ACE2. The dotted line indicates a fold change of 1. Dotted lines within the violins indicate the median and quartiles. N ≥ 3. P < 0.05 *; P<0.01**; P<0.001***; P<0.0001****.

Recently, a novel SARS-CoV-2 variant (clade B.1.1.7 or British variant) emerged and is rapidly replacing endemic viruses globally. Epidemiological data suggest that this variant is 1.35-2 fold more transmissible than the ancestral lineage and is associated with higher viral loads^44–46^. Interestingly, this variant contains a mutation (P681H) directly N-terminal from the RRAR multibasic cleavage site that adds another basic residue to the multibasic cleavage site, creating an HRRAR motif. A similar mutation (P681R) was detected in the Indian variant (clade B.1.617). As the multibasic cleavage site facilitates serine protease-mediated entry^11^, mutations in or near this site may alter protease usage for S2’ cleavage, which directly triggers fusion and entry. We found that the British variant replicated efficiently in wildtype and cathepsin mutant organoids, but not in TMPRSS2-deficient cells (Fig. 4E), indicating that the British variant did not broaden its protease usage. These experiments provide a proof-of-concept on how emerging viral strains could be screened against mutant IOs.

## Discussion

The current COVID-19 pandemic has exposed weaknesses in our preparedness for coronavirus pandemics. No effective coronavirus antivirals are approved for use in humans and all completed large-scale COVID-19 drug trials have failed to show efficacy to this date, including (hydroxy)chloroquine and remdesivir^8,47^. The disappointing clinical effects of (hydroxy)chloroquine in humans in particular highlights gaps in the understanding of fundamental coronavirus biology. (Hydroxy)chloroquine, an inhibitor of the endosomal acidification was identified as a potent inhibitor of SARS-CoV^48^ and SARS-CoV-2^7^ viral entry in cell line-based assay, confirmed here. In agreement with this, recent whole genome CRISPR/Cas9 genetic screens in transformed cell lines again suggested that endosomal entry factors, such as cathepsin L, are crucial for SARS-CoV-2 entry^9,49,50^.

Here, we use human intestinal organoids as a non-transformed model to study genes implicated in coronavirus biology. We have chosen to use only IOs since it is currently not possible to efficiently genetically engineer airway organoids due to limited clonal outgrowth of these cells. Nevertheless, IOs express the majority of host factors assessed, including proteases, to a similar level as the airways (Fig. 1). We confirmed that in this model ACE2 is the obligate entry receptor for SARS-CoV-2 and SARS-CoV, while DPP4 is the entry receptor for MERS-CoV, indicating that accessory receptors may not play crucial roles for these viruses. Indeed, knockout of NRP1, recently proposed as a SARS-CoV-2 (co-)receptor in Hela and Caco-2 cells, did not affect SARS-CoV-2 entry^13,14^. Furthermore, we demonstrate that Cathepsin L and B are not involved in SARS-CoV-2 entry in IOs. In accordance with this, a cathepsin inhibitor (E64D) and chloroquine did not inhibit SARS-CoV-2 in these IOs, while the serine protease inhibitor Camostat effectively blocked viral propagation. A similar anti-SARS-CoV-2 effect of camostat was observed in organoid-derived airway cells^11^. The broad activity of Camostat does not allow to pinpoint which serine protease mediates entry.

TMPRSS2-deficiency in IOs strongly decreased SARS-CoV-2 replication and spread, indicating that TMPRSS2 is the main priming protease. Other related TMPRSS genes have previously also been linked to SARS-CoV-2 replication, including TMPRSS4 in the intestine^33^. Overexpression of TMPRSS11D and TMPRSS13 promoted viral entry into the hamster kidney cell line BHK-21^51^, while TMPRSS4 overexpression facilitated viral entry in HEK293 cells^33^. Like TMPRSS2, TMPRSS4 is highly expressed in intestinal tissue and IOs, yet TMPRSS4 does not appear to rescue loss of TMPRSS2. This discrepancy with previous work may reflect the fact that our study relies on physiological expression of these proteases, rather than on overexpression. Importantly, intestinal organoids express similar TMPRSS family members compared to airway tissue, including high levels of TMPRSS2, TMPRSS3, TMPRSS4 and TMPRSS13. Only the TMPRSS-11A and –11D are relatively enriched in airway-versus intestinal epithelium^52^. Although we cannot exclude the possibility that these TMPRSS family members function in activation of SARS-CoV-2 spike in the airways, their expression levels are much lower than that of TMPRSS2 in airway organoids (Fig. 1A) and lung tissue^52^.

Altogether, these findings indicate that multiple TMPRSS genes may be able to mediate entry when overexpressed, but -at physiological levels in IOs-only TMPRSS2 plays an essential role, which may inspire the development of high-specificity TMPRSS2 inhibitors. The high TMPRSS-2 dependency of SARS-CoV (this study) indicates that such inhibitors may well be effective against future SARS-like coronavirus pandemics. The observation that TMPRSS2-null mice do not display a visible phenotype^53^ implies that such inhibitors may be well-tolerated. Our findings match with observations that SARS-CoV and to a lesser extent MERS-CoV replication and dissemination was reduced in TMPRSS2-deficient mice^54^.

In conclusion, our findings underscore the relevance of non-transformed human models for (corona)virus research and identify TMPRSS2 as an attractive therapeutic target in contrast to many other genes (e.g. cathepsin L, cathepsin B, NRP1, NDST1 etc) that -as deduced from our observations-unlikely to be of clinical value. Future emerging viruses could be readily screened against our IO host factor knockout biobank to rapidly identify therapeutic targets.

### Acknowledgements

We thank Single Cell Discoveries for RNA library preparation, and R. van der Linden and S. van Elst for help with FACS sorting. We thank E. Eenjes and R. Rottier for providing human lung material.

### Author contributions

J.B., M.H.G. and M.M.L. designed the study and performed the experiments. J.P. and J.V. analyzed RNA-sequencing data. B.H. and H.C. supervised the study.

**Supplementary figure 1.**
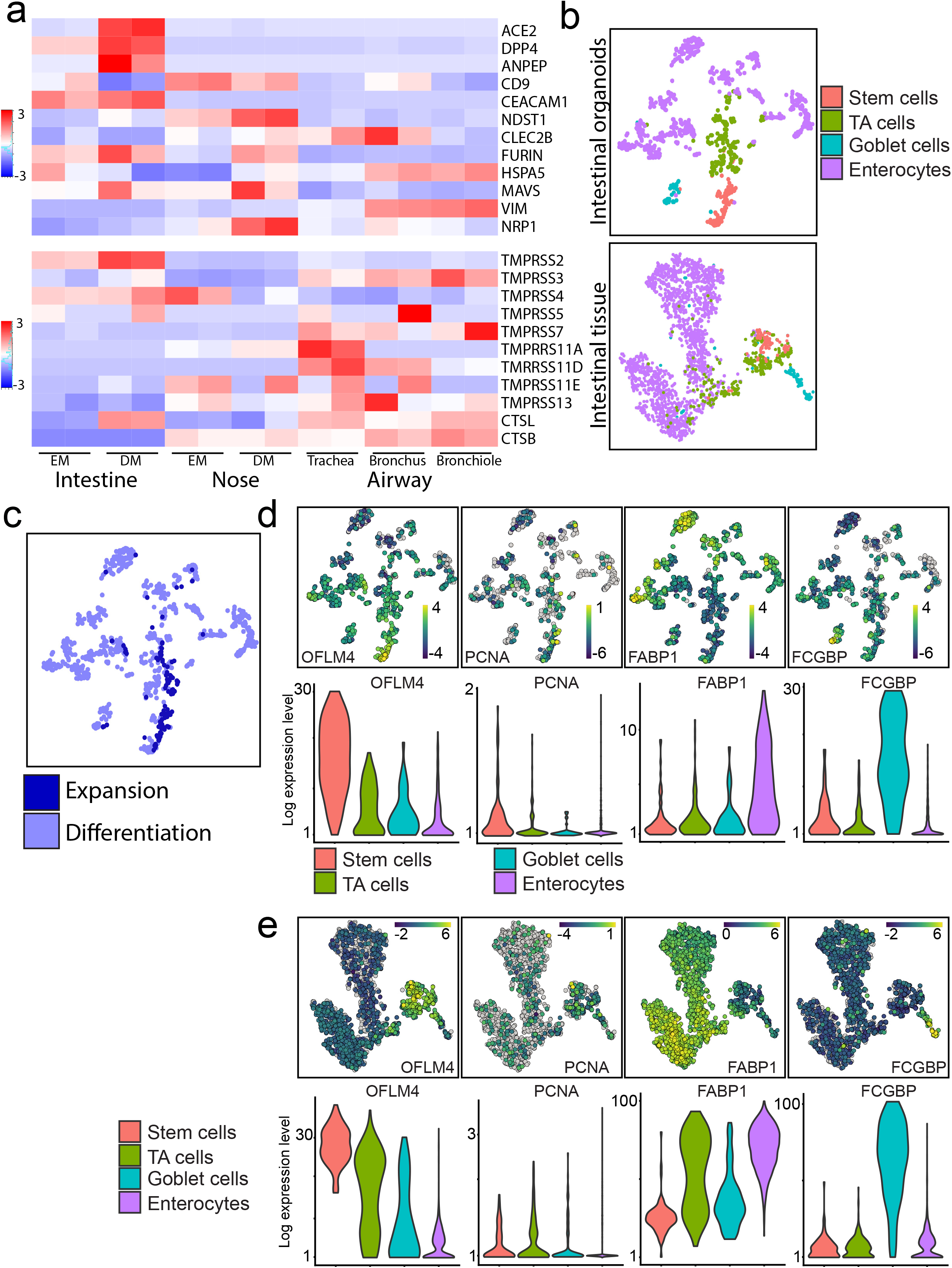
Bulk and single cell RNA sequencing reveals intestinal expression of host proteases involved in viral entry. a) Heatmaps depicting expression of host genes (top) and proteases (bottom) in intestinal, nose and airway organoids. Nose organoids were cultured in expansion (EM) or differentiation (DM) medium. Colored bar represents Z-score of log2 transformed values. b) t-SNE maps displaying a newly generated human organoid single cell RNA sequencing atlas (left), and a dataset reanalyzed from^32^ (right). Colors indicate different cell types. c) t-SNE maps displaying the human organoid single cell sequencing atlas. Color codes indicate cells derived from expansion or differentiation medium. d) t-SNE maps and violin plots displaying expression of host factors in the human intestinal organoid cell atlas. Bars in t-SNE maps display color-coded normalized unique transcript expression (logarithmic scale). e) t-SNE maps and violin plots displaying expression of host factors in the human intestinal tissue atlas. Bars in t-SNE maps display color-coded normalized unique transcript expression (logarithmic scale).

**Supplementary figure 2.**
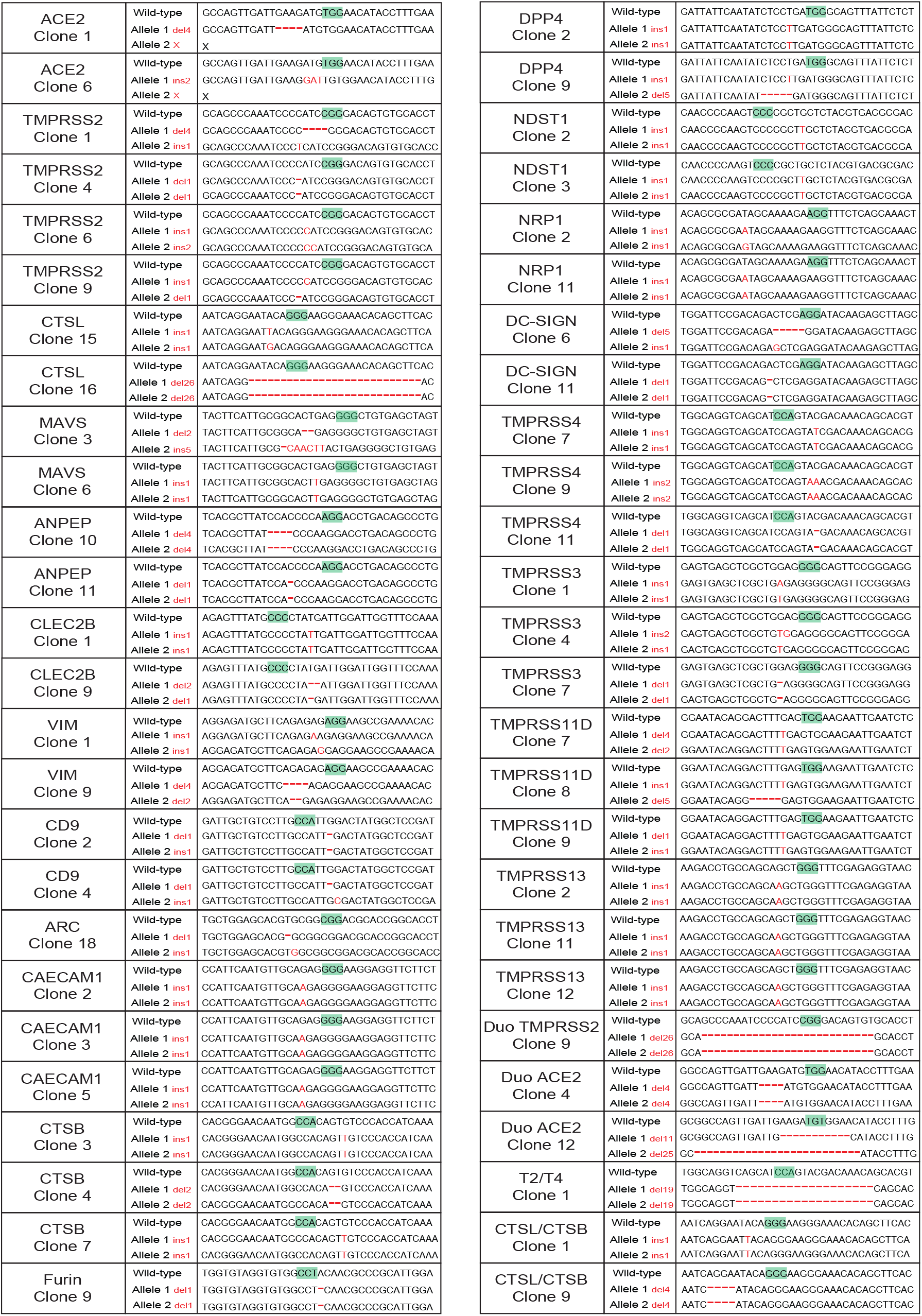
Generation of a host factor loss-of-function organoid biobank. Overview of the genetic alterations causing frameshifts in the different host factors. Green boxes indicate PAM sequence, red dashes or bases indicate respectively deletions and insertions.

**Supplementary figure 3.**
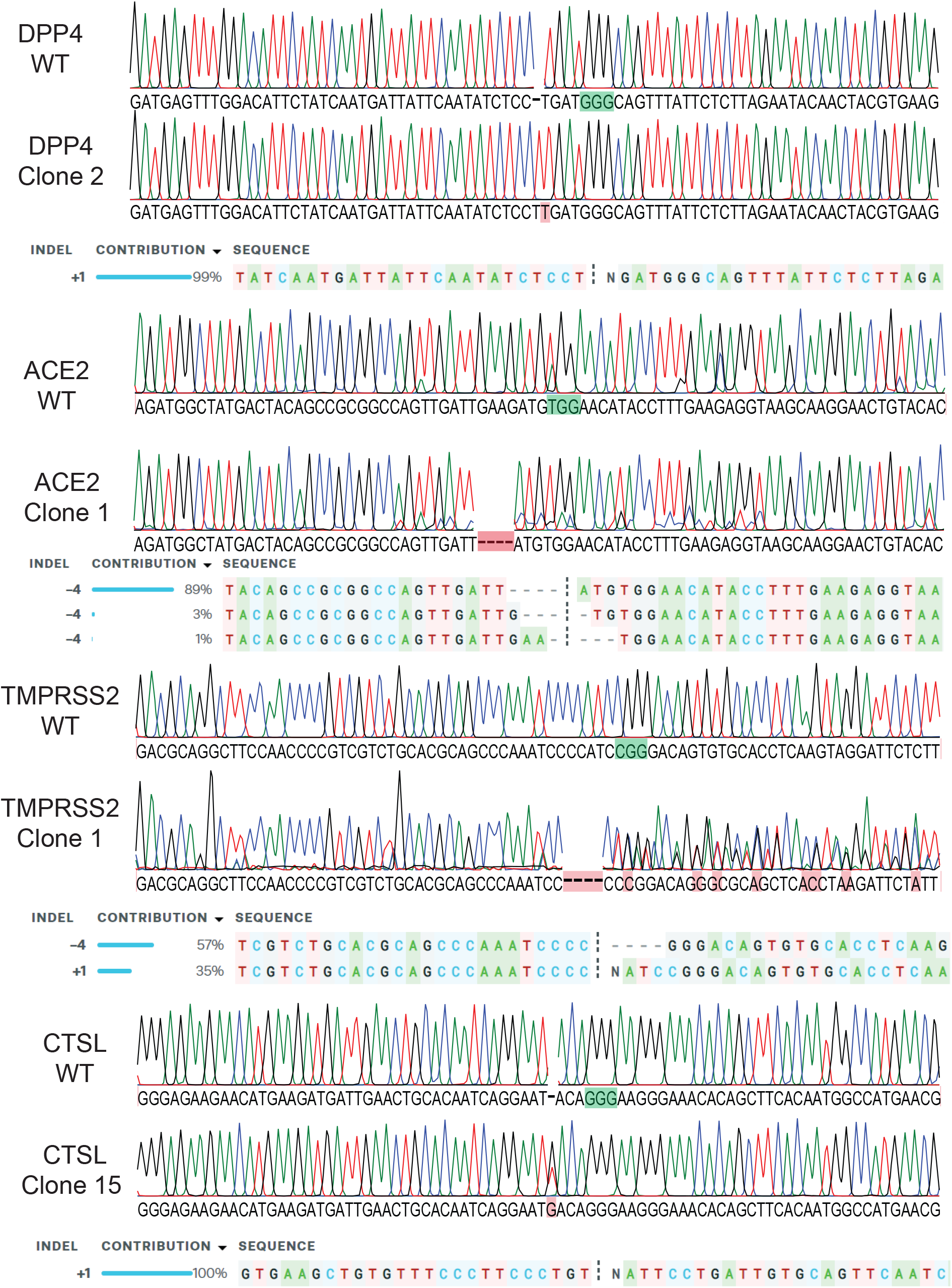
Organoid genotyping by sanger sequencing and *in silico* ICE analysis. Sanger traces and subsequent *in silico* sanger deconvolution by ICE v2 for the first clone of ACE2, TMPRS2, DDP4 and CTSL indicating out-of-frame indel induction at the target site. Green boxes indicate PAM sequence.

**Supplementary figure 4.**
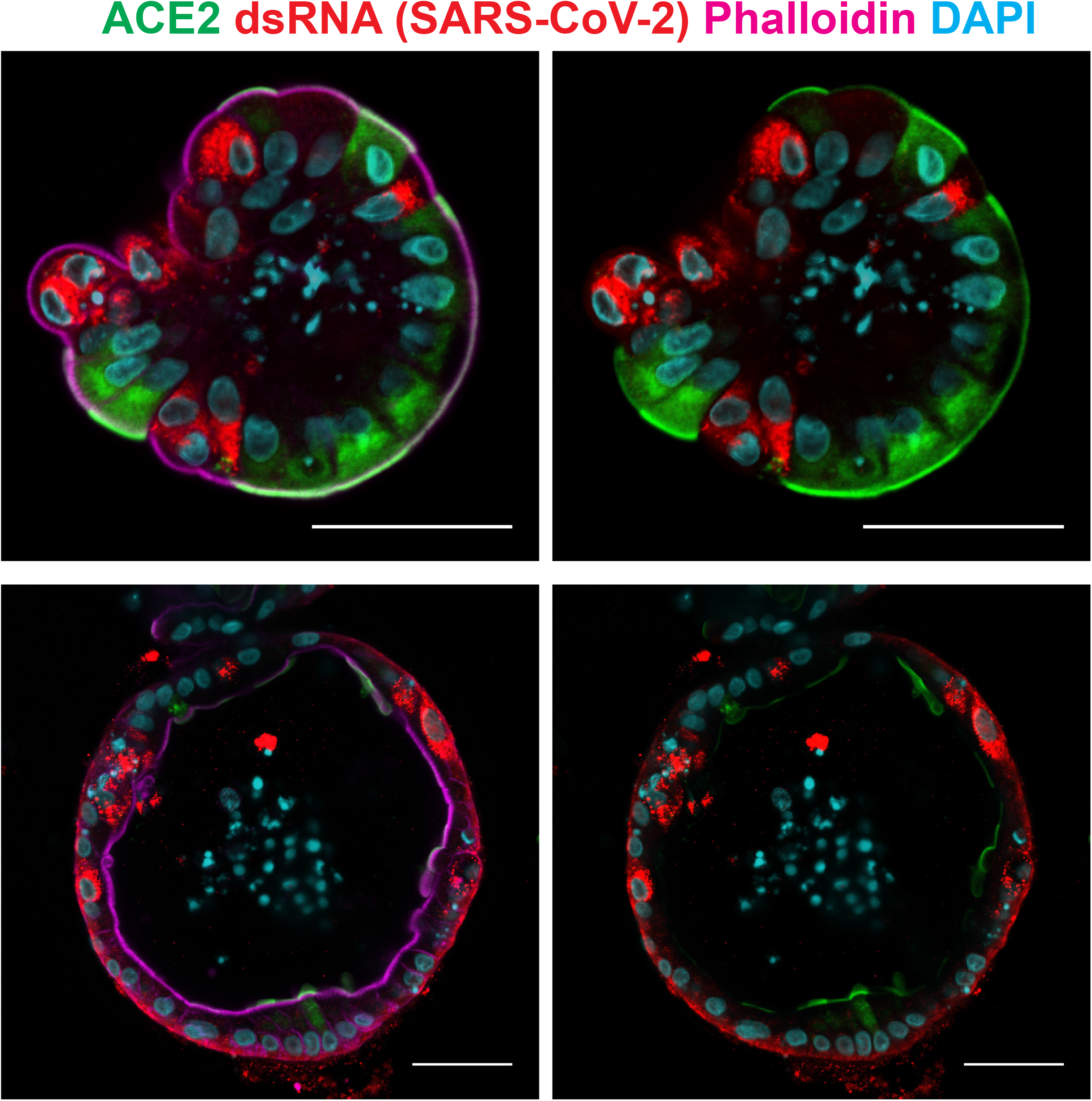
SARS-CoV-2 infected cells contain varying degrees of membranous ACE2 protein. Immunofluorescent staining of SARS-CoV-2 infected organoids. Virus is visualized by dsRNA. Some infected cells are devoid of visible ACE2 on the outer membrane. Scale bars are 50 μm.

**Supplementary figure 5.**
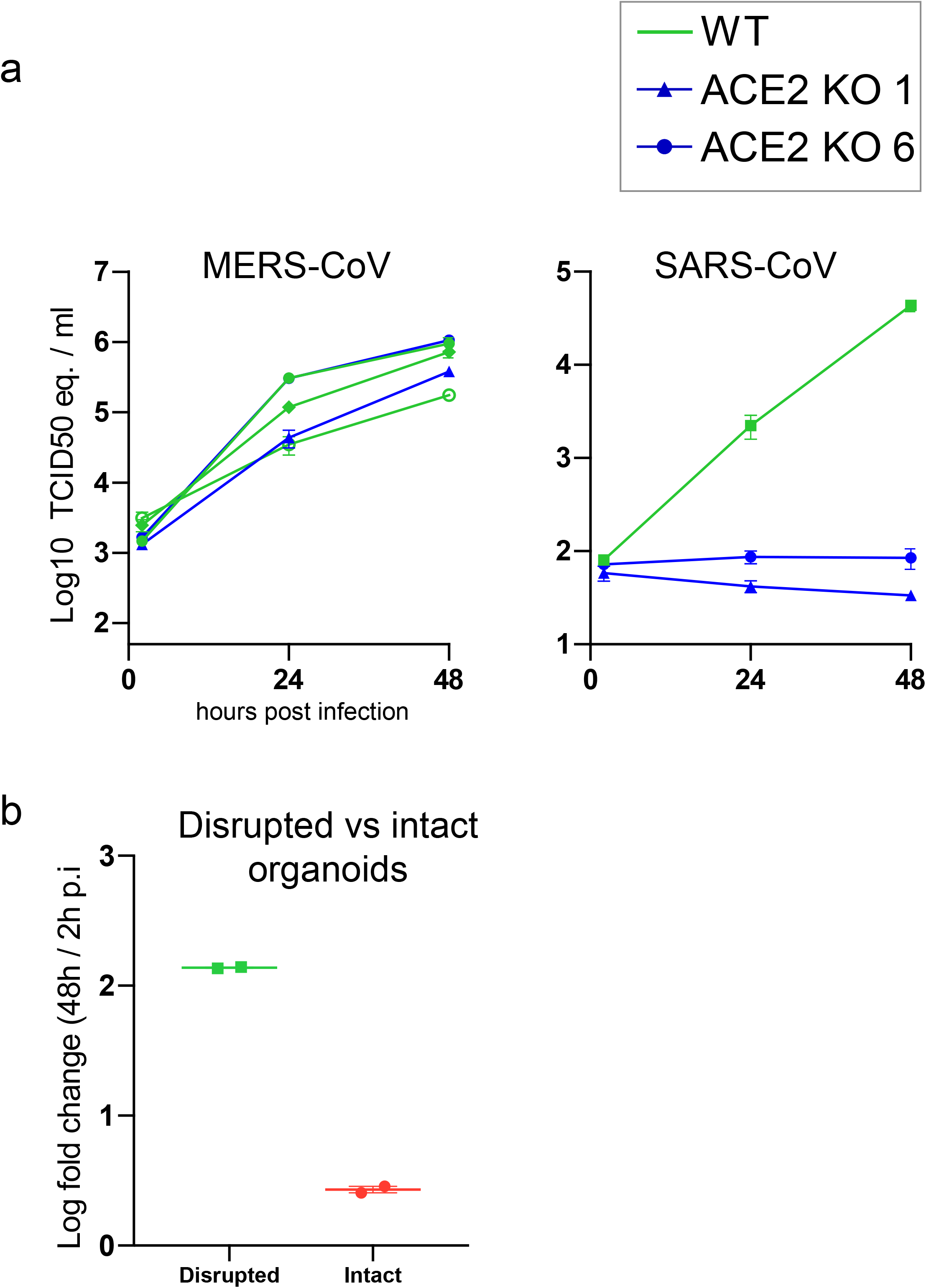
SARS-CoV but not MERS-CoV replication depends on ACE2. a) qPCR analysis targeting the N gene (SARS-CoV) or upE region (MERS-CoV) to quantify viral replication in WT and ACE2 KO organoids. Error bars represent SEM. Each data point represents the mean of 3 replicates. b) qPCR analysis targeting the E gene to quantify viral replication of the SARS-CoV-2 in mechanically disrupted or intact organoids, where virus can only access the basolateral side. Graphs display the ratio between viral titer at 48 hours compared to 2 hours post infection (p.i.). Each data point represents the mean of 3 replicates. Error bars represent SEM.

**Supplementary figure 6.**
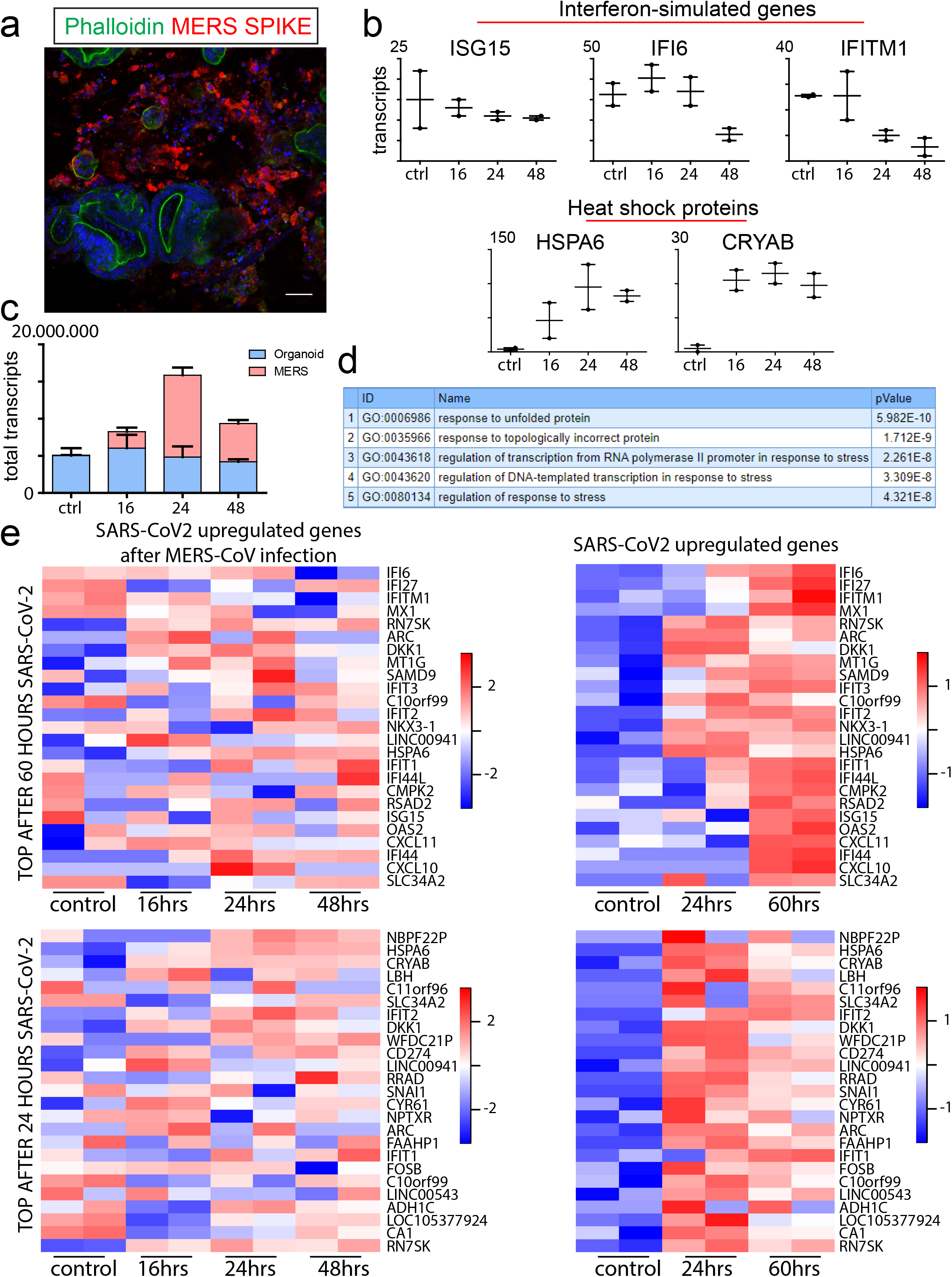
Establishment of MERS infection model in human intestinal organoids. a) Immunofluorescent staining of organoids 48 hours after MERS-CoV infection. Virus is visualized by staining for the spike protein. The majority of the infected organoids display massive cell death. Scale bars are 50 μm. b) Graphs depicting the transcript counts determined by RNA sequencing of different genes upon MERS-CoV infection. Different numbers indicate timepoints (hours) after infection c) Graph depicting the transcript counts mapping to human and MERS genomes in MERS-CoV infected organoids. MERS reads increase over time, but drop again at 48 hours potentially due to cell death of infected cells. For all other analyses, MERS reads were removed from analyses for normalization purposes. d) Go term enrichment analysis for biological processes of the 60 most significantly upregulated genes upon MERS-CoV infection in organoids. e) Heatmap depicting the expression profile of the 25 genes with strongest upregulation upon SARS-CoV-2 infection^17^; right heatmaps) and the same genes upon MERS-CoV infection (left heatmaps). The top heatmaps show the most prominently upregulated genes after 60 hours of SARS-CoV-2 infection, the lower heatmaps after 24 hours. In contrast to SARS-CoV-2, MERS-CoV does not induce expression of ISGs. Colored bar represent Z-score of log2 transformed values.

**Supplementary figure 7.**
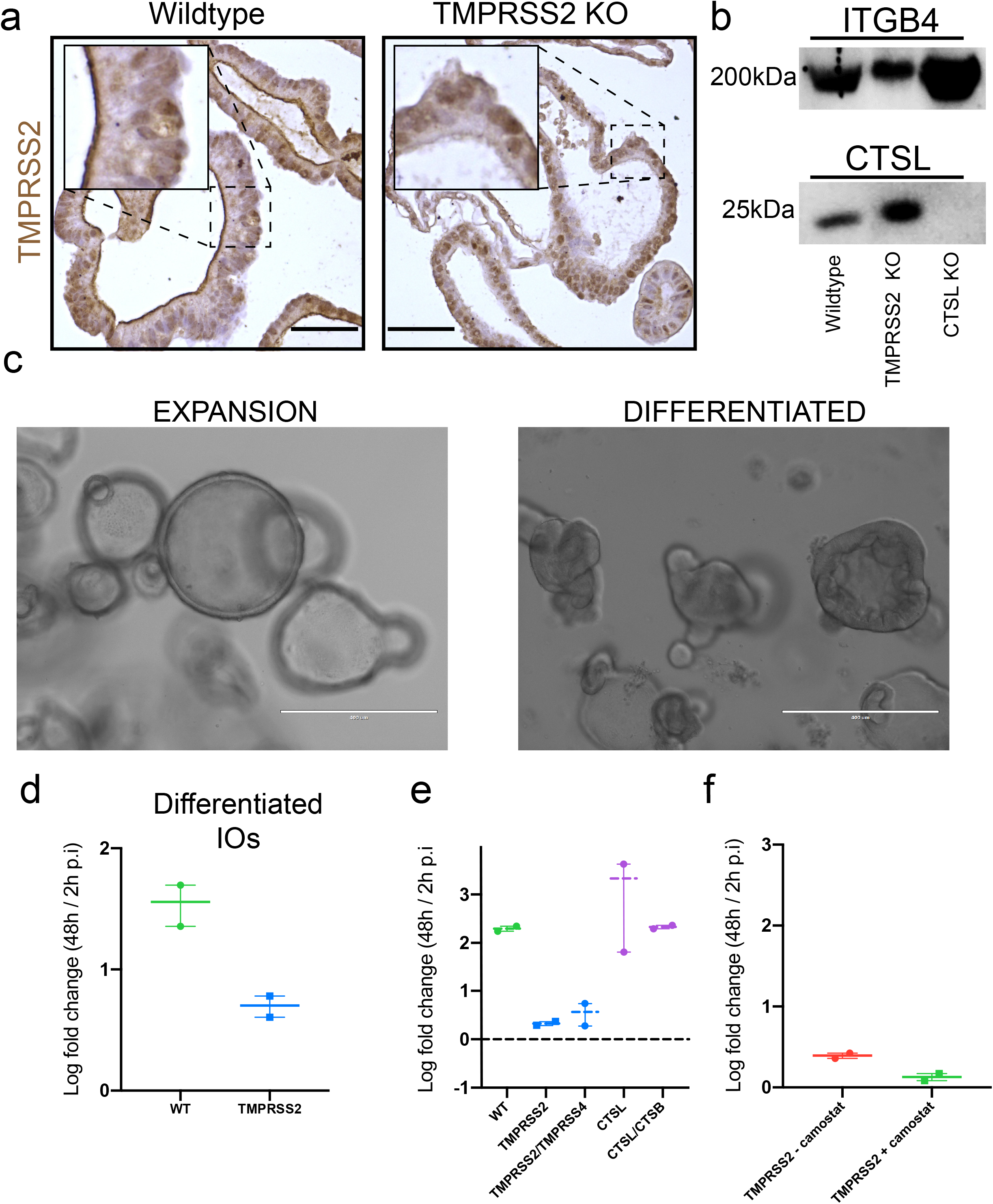
Lack of redundancy in cathepsins and serine proteases in viral entry. a) Immunohistochemical staining of TMPRSS2 in wildtype and TMPRSS2-knock out (KO) organoids. TMPRSS2 locates mostly to the apical membrane in wildtype cells, and is absent in mutant organoids. Scale bars are 50 μm. b) Western blotting for CTSL and Integrin B4 (ITGB4, loading control) in wildtype, TMPRSS2- and CTSL-KO organoids. CTSL protein is completely lost in corresponding mutant organoids c) Brightfield images of expanding and 5-day differentiated organoids that were infected with SARS-CoV-2. Scale bars are 400 μm. d) Graph displaying the ratio between the viral titer at 48 hours compared to 2 hours post infection (p.i.) quantified by qPCR targeting the E gene to measure viral replication of SARS-CoV-2 in expanding and 5-day differentiated intestinal organoid cells harboring a loss-of-function mutation in the TMPRSS2 gene. Error bars represent SEM. N=2. e) qPCR analysis targeting the E gene to quantify viral replication of the SARS-CoV-2 in organoids harboring different single and double mutants in host proteases. Graphs display the ratio between viral titer at 48 hours compared to 2 hours post infection (p.i.). Error bars represent SEM. The dotted line indicates a fold change of 1. N=2. f) qPCR analysis targeting the E gene to quantify viral replication of the SARS-CoV-2 in TMPRSS2-deficient organoids treated with the broad serine protease inhibitor Camostat. Graphs display the ratio between viral titer at 48 hours compared to 2 hours post infection (p.i.). Error bars represent SEM. N=2.

**Supplementary figure 8.**
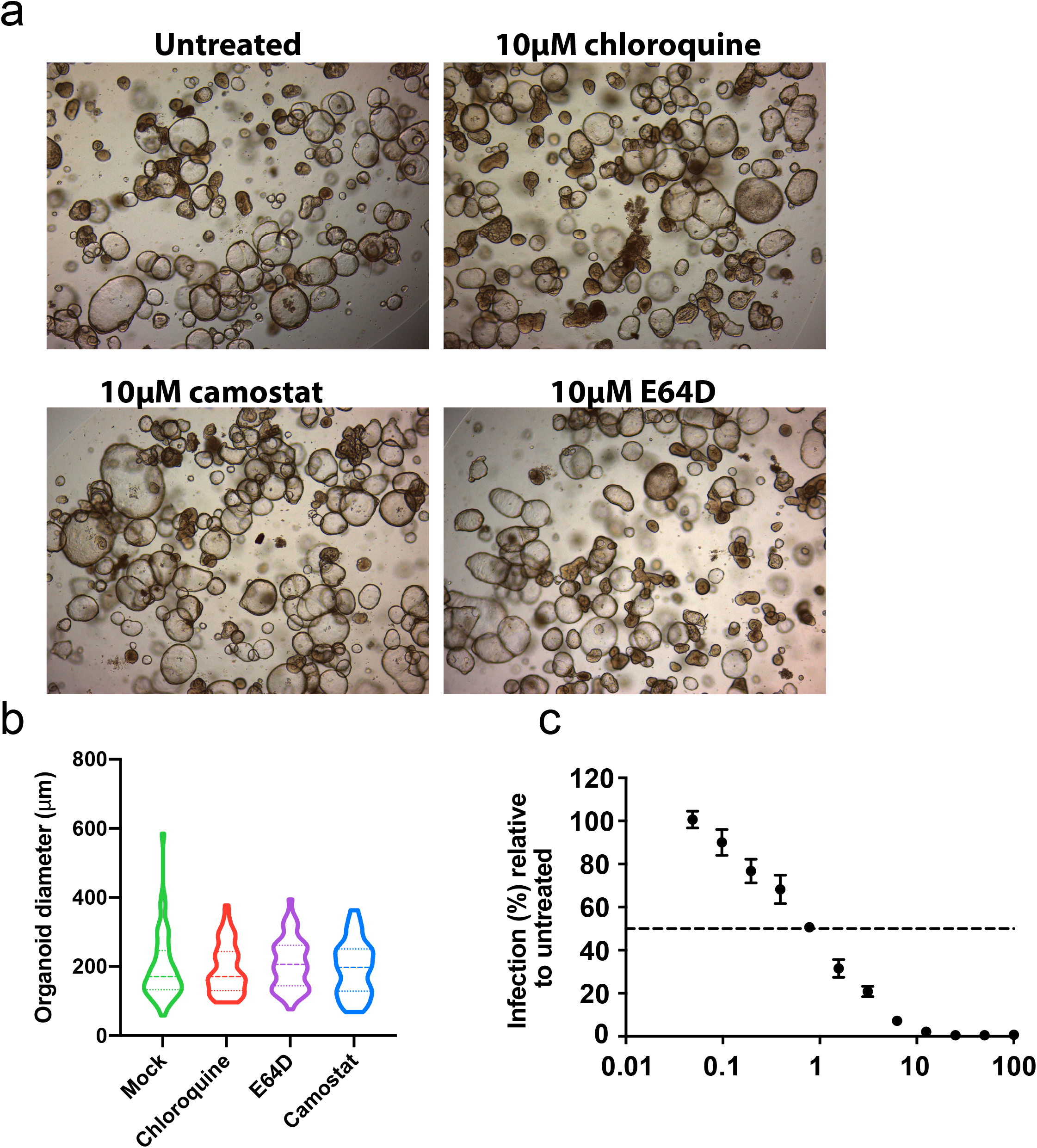
Inhibition of serine proteases but not chloroquine inhibits viral replication in organoids. a) Brightfield images of organoids treated for 48 hours with serine protease inhibitor Camostat, chloroquine or cysteine protease inhibitor E64D. b) Violin plot of average sizes in organoids from Fig. 6E. The diameter was measured in at least n=50 organoids per treatment. Organoid size was not significantly changed in any of the treatments, indicating similar growth. Dotted lines within the violins indicate the median and quartiles. c) Quantification of viral entry in Vero E6 cells upon treatment with chloroquine using immunostaining 8 hours after infection. Error bars represent SEM.

**Supplementary figure 9.**
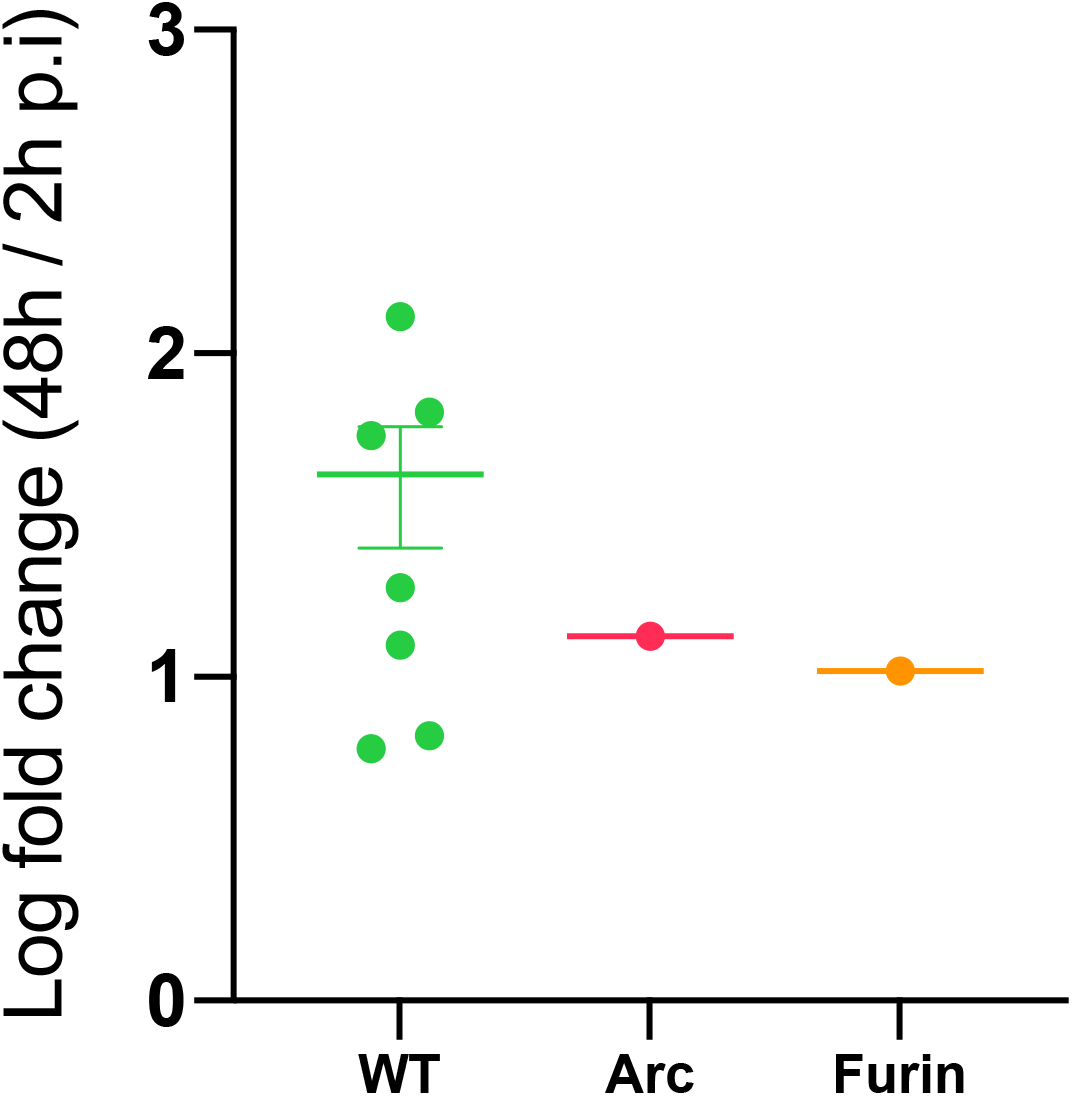
SARS-CoV-2 replication in Furin-en ARC-mutant IOs. Graph displaying the ratio between the viral titer at 48 hours compared to 2 hours post infection (p.i.) quantified by qPCR targeting the E gene to measure viral replication of SARS-CoV-2 in wildtype, and ARC- and Furin-mutant organoids. Experiment was performed with n=1 biological replicate. The WT data is redispayed from Fig. 4A.

**Supplementary table 1 Normalized transcript counts in intestinal and airway organoids**

Table shows normalized transcript counts determined by RNA sequencing of duplicate organoids from the respective regions, and intestinal organoids infected with SARS-CoV-2. The intestinal organoid dataset was obtained from^17^.

**Supplementary table 2 Overview of genetically modified organoids generated in this study**

**Supplementary table 3 Normalized transcript counts in MERS-CoV infected organoids**

Table shows normalized transcript counts determined by RNA sequencing of a duplicate control treatment (NC), and 16, 24 and 48 hours after MERS-CoV infection.

**Supplementary table 4 Differentially regulated genes upon MERS-CoV infection in organoids**

Fold change in gene expression versus control after 48 hours of MERS-CoV infection.

**Supplementary table 5 Oligos used in this study as gRNAs and sequencing primers**

## Methods

### Cell culture of human intestinal organoids and human airway

Human small intestinal tissue was obtained from the UMC Utrecht with informed consent of the patient. The patient was operated for a colorectal tumor, and a sample from non-transformed, normal mucosa was taken for this study. The study was approved by the UMC Utrecht (Utrecht, The Netherlands) ethical committee and was in accordance with the Declaration of Helsinki and according to Dutch law. This study is compliant with all relevant ethical regulations regarding research involving human participants.

Nasal inferior turbinate brushes were obtained from the Hadassah Medical Center, Jerusalem, with informed consent of the patient. Patients were diagnosed with primary ciliary dyskinesia, and tissue was obtained from healthy donors as a comparison. Healthy material was used for the RNA sequencing in this work. The study was approved by the ethical committee and was in accordance with the Declaration of Helsinki and according to Israeli law under IRB approval number 075-16 HMO. This study is compliant with all relevant ethical regulations regarding research involving human participants.

Adult lung tissue was obtained from residual, tumor-free, material obtained at lung resection surgery for lung cancer. The Medical Ethical Committee of the Erasmus MC Rotterdam granted permission for this study (METC 2012-512).

Human small intestinal cells were isolated, processed and cultured as described previously^55,56^. Wnt surrogate was used (0,15nM, U-Protein Express) instead of Wnt conditioned media. Differentiation of intestinal organoids was achieved as described previously^56^.

Nose tissue was dissociated and cultured as described previously^57^. Differentiation towards ciliated cells was performed by activating BMP signaling and inhibiting Notch signaling for 10 days (Van der Vaart et al., under review)

Isolation of human bronchial airway stem cells was performed using a protocol similar to Sachs and colleagues^57^. Small airway stem cells were isolated from distal human lung parenchyma as described before^57^. Tracheal stem cells were collected from tracheal aspirates of intubated preterm infants (28 weeks gestational age). Organoids were cultured as described before^57^. To obtain differentiated organoid-derived cultures, organoids were dissociated into single cells using TrypLE express (Gibco; #12604013). Cells were seeded on Transwell membranes (Corning) coated with rat tail collagen type I (Fisher Scientific). Single cells were seeded in AO growth medium: complete base medium (CBM; Stemcell Pneumacult-ALI; #05001) at a 1:1 ratio. After 2-4 days, confluent monolayers were cultured at air‐liquid interphase in CBM. Medium was changed every 5 days for 8 weeks.

### Transfection of organoids for CRISPR-Cas9 experiments

sgRNAs targeting loci of interest were cloned into a SpCas9-EGFP vector (addgene plasmid #48138) using a protocol described before^58^. sgRNAs were designed using WTSI website (https://www.sanger.ac.uk/htgt/wge/). A full list of gRNAs and primers to generate SpCas9-EGFP expressing plasmids can be found in supplementary table 5. To generate homozygous frameshift mutations in genes of interest, organoids were transfected with SpCas9-EGFP containing the locus-specific sgRNA. Transient transfection using a NEPA21 electroporator was performed as described before^59^. 3-7 days after transfection, organoids were dissociated using TryplE (TryplE Express; Life Technologies) and sorted on a FACS-ARIA (BD Biosciences) for GFP positivity. After sorting, Rho kinase inhibitor (Y-27632 dihydrochloride; 10μM, Abmole) was added for 1 week to support single cell outgrowth.

### Generation of stable genetically modified organoid lines

To generate clonal organoid lines with genotypes of preference, organoids were picked 2 weeks after sorting. Manually picked organoids were dissociated using TryplE (TryplE Express; Life Technologies) and plated in BME in pre-warmed cell culture plates. After two weeks, single cells grew into organoids and were split again to verify actively dividing stem cells. After the second split, 20μL of organoid-BME suspension was directly taken from the plate and DNA was extracted from the organoids using the Zymogen Quick-DNA microprep kit according to protocol. Regions around sgRNA target sites were amplified using Q5 high fidelity polymerase (NEB) according to manufacturer’s protocol. CRISPR/Cas9-mediated indel formation was confirmed by sanger sequencing of these amplicons (Macrogen). Subsequently, sanger trace deconvolution was performed with the use of ICE v2 CRISPR analysis tool (synthego website) to call clonal organoid lines with homozygous frameshift mutations at the target site. Knockout clones were further expanded for viral infection experiments. Primers used for amplification and sanger sequencing can be found in supplementary table 5. For the generation of TMPRSS2/TMPRSS4 double mutants, TMPRSS4 was knocked out in TMPRSS2-clone 9. For the generation of CTSL/CTSB double mutants CTSL was knocked out in CTSB clone 3.

### Viruses and cell lines

Vero and VeroE6 cells were maintained in Dulbecco’s modified Eagle’s medium (DMEM, Gibco) supplemented with 10% fetal calf serum (FCS), HEPES, sodium bicabonate, penicillin (100 IU/mL) and streptomycin (100 IU/mL) at 37°C in a humidified CO2 incubator. Calu-3 cells were maintained in Opti-MEM I (1X) + GlutaMAX (Gibco)(Gibco) supplemented with 10% FCS, penicillin (100 IU/mL) and streptomycin (100 IU/mL) at 37°C in a humidified CO2 incubator. SARS-CoV (isolate HKU 39849, genbank accession no. AY278491), SARS-CoV-2 (isolate Bavpat-1; European Virus Archive Global #026V-03883; kindly provided by Dr. C. Drosten) were propagated on VeroE6 cells in Opti-MEM I (1X) + GlutaMAX (Gibco), supplemented with penicillin (100 IU/mL) and streptomycin (100 IU/mL) at 37°C in a humidified CO2 incubator. MERS-CoV (isolate EMC, genbank accession no. NC019843) was propagated on Vero cells in the same medium. Non-adapted SARS-CoV-2 Bavpat-1 and B.1.1.7 (genbank accession no. MW947280) were propagated in Calu-3 cells as described before^39^. Stocks were produced by infecting cells at a multiplicity of infection (MOI) of 0.01 and incubating the cells for 72 hours. The culture supernatant was cleared by centrifugation and stored in aliquots at −80°C. Stock titers were determined by preparing 10-fold serial dilutions in Opti-MEM I (1X) + GlutaMAX. Aliquots of each dilution were added to monolayers of 2 × 104 VeroE6 (for SARS-CoV and SARS-CoV-2) or Vero cells (for MERS-CoV) in the same medium in a 96-well plate. Plates were incubated at 37°C for 5 days and then examined for cytopathic effect. The TCID50 was calculated according to the method of Spearman & Kärber. All work with infectious SARS-CoV, SARS-CoV-2 and MERS-CoV was performed in a Class II Biosafety Cabinet under BSL-3 conditions at Erasmus Medical Center.

### SARS-CoV, SARS-CoV-2 and MERS-CoV infection

Infections were performed using a protocol similar to^17^. Briefly, organoids were harvested in cold Advanced DMEM (including HEPES, Glutamax and antibiotics, termed AdDF+++^17^), washed once in AdDF+++, and sheared using a flamed Pasteur pipette in AdDF+++. Differentiated organoids were broken using a 5-minute incubation with TrypLE (TrypLE Express; Life Technologies). After shearing, organoids were washed once in AdDF+++ before infection was performed in expansion medium. For the experiment in Figure S5B, organoids were gently harvested using a wide pipet tip to avoid shearing organoids. A multiplicity of infection (MOI) of 0.1 was used for SARS-CoV and SARS-CoV-2 and an MOI of 0.2 was used for MERS-CoV. After 2 hours of virus adsorption at 37°C 5% CO2, cultures were washed four times with 4 ml AdDF+++ to remove unbound virus. Organoids were re-embedded into 30 μL BME in 48-well tissue culture plates and cultured in 250 μL expansion or differentiation medium at 37°C with 5% CO2. Each well contained ~200,000 cells per well. At indicated time points cells were harvested by resuspending the BME droplet containing organoids into 200 μL AdDF+++. Samples were stored at −80°C, a process which lysed the organoids, releasing their contents into the medium upon thawing.

For testing the antiviral activity of chloroquine diphosphate (Sigma), camostat mesylate (Sigma) and E64D (Medchemexpress) in intestinal organoids, sheared organoids were preincubated with these compounds in AdDF+++ at the indicated concentrations for 30 min at 37°C 5% CO2 before infection at an MOI of 0.1 with SARS-CoV-2. After virus adsorption for 2 hours at 37°C 5% CO2, organoids were washed and re-embedded in BME as indicated above. Medium containing the inhibitors was added to the wells for the duration of the experiment. Cells were harvested at indicated time points as described above and stored at −80°C.

### SARS-CoV-2 entry inhibition assay by chloroquine in VeroE6 cells

Chloroquine was two-fold serially diluted in Opti-MEM I (1X) + GlutaMAX starting from a concentration of 100 μg/mL. 100 μl of each dilution was added to a confluent 96-well plate of VeroE6 cells and pre-incubated at 37°C 5% CO2 for 30 minutes. Next, cells were incubated with 400 plaque-forming units of virus in the same concentration range of chloroquine at 37°C 5% CO2. After 8 hours incubation, cells were fixed with formalin, permeabilized with 70% ethanol and stained with polyclonal rabbit anti-SARS-CoV nucleoprotein antibody (1:1000; 40588-T62, Sino Biological) followed by secondary Alexa488 conjugated goat-anti-rabbit antibody (Invitrogen). Plates were scanned on the Amersham™ Typhoon™ Biomolecular Imager (channel Cy2; resolution 10μm; GE Healthcare). Data was analyzed using ImageQuant TL 8.2 image analysis software (GE Healthcare).

### Determination of virus titer using qRT-PCR

For determining the viral titer using qPCR, samples were thawed and centrifuged at 2,000 g for 5 min. Sixty μL supernatant was lysed in 90 μL MagnaPure LC Lysis buffer (Roche) at RT for 10 min. RNA was extracted by incubating samples with 50 μL Agencourt AMPure XP beads (Beckman Coulter) for 15 min at RT, washing beads twice with 70% ethanol on a DynaMag-96 magnet (Invitrogen) and eluting in 30 μL MagnaPure LC elution buffer (Roche). Viral titers (TCID50 equivalents per mL) were determined by qRT-PCR using primer-probe sets described previously^60–62^ and comparing the Ct values to a standard curve derived from a titrated virus stock.

### Immunostainings and western blot

Organoids were stained as described before^55^. Whole organoids were collected by gently dissolving the BME in ice-cold PBS, and subsequently fixed overnight at 4°C in 4% paraformaldehyde (Sigma). Next, organoids were permeabilized and blocked in PBS containing 0,5% Triton X-100 (Sigma) and 2% normal donkey serum (Jackson ImunoResearch) for 30 min at room temperature. All stainings were performed in blocking buffer (2% normal donkey serum in PBS). For immunofluorescence, primary antibodies used were mouse anti-nucleoprotein (1:200; 40143-MM05, Sino Biological), mouse anti-dsRNA (1:200; Scicons), goat anti-ACE (1:100; R&D Systems, AF933), goat anti-DPP-4 (1:200; R&D systems, AF1180) and rabbit anti-MERS S1 (1:200; 40069-T52, Sino Biological). For immunofluorescence, organoids were incubated with the corresponding secondary antibodies Alexa488-, 568- and 647-conjugated anti-rabbit and anti-goat (1:1,000; Molecular Probes) or Phalloidin-Alexa488 (Thermofisher Scientific) in blocking buffer containing 4ʹ,6-diamidino-2-phenylindole (DAPI; 1;1,000, Invitrogen). After staining, organoids were transfected to a glass slide embedded in Vectashield (Vector labs). Stained organoids were imaged using a SP8 confocal microscope (Leica) or a Zeiss LSM700, and image analysis and presentation was performed using ImageJ software.

Immunohistochemistry was performed as described before^63^. Antigen retrieval for TMPRSS2 staining was achieved by boiling for 20 minutes in citrate buffer. Primary antibody used was rabbit anti-TMPRSS2 (1:100; Abcam, ab109131) followed by anti-rabbit conjugated to horseradish peroxidase (Powervision, Leica)

For Western blot of CTSL, organoid proteins were solubilized using a standard RIPA buffer for 30 min on ice in the presence of protease inhibitors. Samples were run on a 4-15% PAA gel (BioRad) under reducing conditions. Proteins were electrophoresed to PVDF membranes from Immobilon.Both primary antibodies, mouse anti-CTSL (± 25 kDa) and mouse anti-ITGB4 (± 200 kDa), were incubated O/N at 4C in PBS/10% milk protein/0.1% Tween20. The secondary rabbit anti-mouse HRP-conjugate (Dako) was incubated for 2hrs at 4C in the same buffer. The mouse IgG2a antibody against ITGB4, 58XB4, was a gift from A. Sonnenberg (NKI, Amsterdam, The Netherlands). HRP activity was visualized with ECL (GE-Healthcare).

### Bulk RNA sequencing

Single Cell Discoveries (Utrecht, The Netherlands) performed library preparation, using an adapted version of the CEL-seq protocol, as we have done previously^17^. After library generation, paired-end sequencing was performed on the Illumina Nextseq500 platform using barcoded 1 × 75 nt read setup. Read 1 was used to identify the Illumina library index and CEL-Seq sample barcode. Read 2 was aligned to the CRCh38 human RefSeq transcriptome, with the addition SARS-CoV-2 (Ref-SKU: 026V-03883) or MERS (NC_038294.1) genomes, using BWA using standard settings^64^. Reads that mapped equally well to multiple locations were discarded. Mapping and generation of count tables was performed using the in-house MapAndGo script, filtering to exclude reads with identical library- and molecule-barcodes. RNA sequencing data from expanding and differentiated human intestinal organoids, infected with SARS-CoV-2, was used from a previous publication^17^. Normalization using the median of ratios method and differential gene expression analysis was performed using the DESeq2 package^64^. SARS- and MERS-mapping reads were removed before normalization to avoid biasing organoid transcript counts. To generate heatmaps, row z-scores of selected genes were calculated from the samples selected.

### Organoid preparation for single cell sequencing analysis

Human ileal organoids were differentiated as previously described^56^. A control condition was kept inhuman organoid expansion medium to obtain stem- and progenitor cells for comparison.

Dissociation of organoids to single cells was performed by a 10-minute incubation with TrypLE (TrypLE Express; Life Technologies) supported by repeated mechanical disruption using a narrowed glass pipette. Viable cells were sorted using a BD FACS Aria (BD Biosciences) using DAPI exclusion. Individual cells were collected in 384-well plates with ERCC spike-ins (Agilent), reverse transcription primers and dNTPs (both Promega). Single cell sequencing was performed according to the Sort-seq method^65^.

Sequencing libraries were generated with TruSeq small RNA primers (Illumina) and sequenced paired-end at 60 and 26 bp read length, respectively, on the Illumina NextSeq.

### Single cell RNA sequencing analysis from intestinal organoids and tissue

Reads derived from 1344 cells (192 expansion medium, 1152 differentiation medium) were mapped to the human GRCh37 genome assembly. Sort-seq read counts were filtered to exclude reads with identical library-, cell- and molecule barcodes. UMI counts were adjusted using Poisson counting statistics^65^. Cells with fewer than 1,000 unique transcripts were excluded from further analysis. This resulted in 944 remaining cells (126 from expansion, 818 from differentiation medium)

Subsequently, RaceID3 was used for k-medoids-based clustering (knn = 5; cln = 20) of cells and differential gene expression analysis between clusters using the standard settings described at https://github.com/dgrun/RaceID3_StemID2_package. Cell types were annotated by cluster based on the expression of marker genes (OLFM4 for stem cells, FABP1 for enterocytes, FCGBP for goblet cells. Lack of these and expression of cell cycle markers including PCNA defined proliferating progenitors cells).

For comparison with tissue-derived cells, we reanalyzed a previously published dataset^32^ of primary human ileal cell types. To compare with the organoid data set, cells were required to be annotated as stem cell, progenitor cell, goblet cell or enterocyte and exhibit more than 3,000 unique transcripts. This resulted in 2137 included cells, which were subsequently clustered using the standard settings of RaceID3 (cln = 16). Cells were assigned an identity based on their annotation from^32^.

### Quantification and statistics

No statistical methods were used to predetermine sample size. The experiments were not randomized and the investigators were not blinded to the sample allocation during experiments and outcome assessment. All data are presented as mean ± standard error of the mean (SEM), unless stated otherwise. Value of n is always displayed in the figure as individual data points, and in the legends.

Statistical analysis was performed with the GraphPad Prism 9 software. We compared differences in virus replication and organoid growth by one-way ANOVA followed by a multiple-comparison test (Original FDR method of Benjamini and Hochberg; Q = 0.05) on log10 transformed values. Statistics were applied if N ≥ 3.

## Data availability statement

All bulk and single cell RNA sequencing data of this study has been uploaded to the Gene Expression Omnibus (GEO), and will be publically available upon publication (GEO accession number is pending).

